# The Plasmodium falciparum NCR1 membrane protein is a novel antimalarial target that exports cholesterol to maintain membrane homeostasis

**DOI:** 10.1101/2024.09.30.615907

**Authors:** Zhemin Zhang, Meinan Lyu, Xu Han, Sepalika Bandara, Meng Cui, Eva S. Istvan, Xinran Geng, Marios L. Tringides, William D. Gregor, Masaru Miyagi, Jenna Oberstaller, John H. Adams, Youwei Zhang, Marvin T. Nieman, Johannes von Lintig, Daniel E. Goldberg, Edward W. Yu

## Abstract

Malaria is an extremely devastating parasitic infection that kills over half a million people each year. It is the leading cause of death in many developing countries, in part, due to a lack of resources and readily available therapeutics. Unfortunately, the most prevalent and deadliest causative agent of malaria, *Plasmodium falciparum*, has developed resistance to nearly all currently available antimalarial drugs. The *P. falciparum* Niemann-Pick Type C1-related (PfNCR1) transporter has been identified as a druggable target, as it is required for membrane homeostasis of the parasite. However, the structure and detailed molecular mechanism of this membrane protein are not yet available. Here we present three structures of PfNCR1 both in the absence and presence of the functional inhibitor MMV009108 at resolutions between 2.98 Å and 3.81 Å using single-particle cryo-electron microscopy (cryo-EM). The data suggest that PfNCR1 binds cholesterol and forms a cholesterol transport tunnel to modulate the composition of the parasite plasma membrane. Cholesterol efflux assays substantiate this as they show that PfNCR1 is an exporter capable of extruding cholesterol from the membrane. Additionally, the inhibition mechanism of MMV009108 appears to be due to a direct blockage of PfNCR1, preventing this transporter from shuttling cholesterol.

## Introduction

Malaria remains a persistent global health problem with the most vulnerable groups being young children and pregnant women. *Plasmodium falciparum* is the most prevalent and deadliest causative agent of malaria. In 2021, this parasite was responsible for causing the disease in approximately 247 million people and claiming 619,000 lives, with 96% of the mortalities located in Africa (*1*). Unfortunately, *P. falciparum* has developed resistance to nearly all currently available antimalarial drugs, including sulfadoxine, pyrimethamine, mefloquine, halofantrine and quinine (*2*). Resistance to highly effective artemisinin-combination therapies first emerged in parts of southeast Asia (*3*) and may soon migrate to Africa. The impact of multidrug resistant malaria is an enormous global health threat, and the identification of novel classes of antimalarials and druggable targets should be a global health priority.

It is known that the malarial parasite *P. falciparum* relies on cholesterol (Chl) for growth and invasion. However, *P. falciparum* is not capable of synthesizing this essential molecule *de novo* and must acquire Chl from the host for survival and development (*4*). Indeed, *P. falciparum* expresses several proteins that likely play important roles in uptake, binding and intracellular trafficking of Chl (*5*). Intriguingly, it has been observed that several human host cells, including liver and red blood cells, are rich in Chl, and these cells are particularly favorable for malarial parasites to reside and grow (*5*).

Although Chl is a prerequisite to the invasion and growth of *P. falciparum* in the host, it appears that this can be a double-edged sword, as the parasite has to carefully control and maintain a certain level of Chl for successful infection. It has been observed that artificially reducing the Chl concentration in erythrocytes leads to less infected cells (*6–8*). However, artificially increasing the Chl level to high concentrations in erythrocytes results in a decrease in the efficiency of the parasite’s invasion (*6, 9*). After invasion, the parasite must maintain the level of Chl in the host’s plasma membrane above a certain threshold. It has been observed that the removal of Chl from the erythrocyte plasma membrane leads to a rapid extrusion of the parasite from the infected erythrocyte (*8*). On the other hand, artificially enhancing the accumulation of Chl within the parasites can lead to death in the infected cell (*10*). Therefore, *P. falciparum* must develop an effective mechanism to regulate Chl level and membrane homeostasis.

*P. falciparum* has a large number of membrane proteins with unidentified function. One such membrane protein is the *P. falciparum* Niemann-Pick Type C1-Related protein (PfNCR1), which is encoded by the gene PF3d7_0107500 (*11*). PfNCR1 is homologous to the human Niemann-Pick C1 (hNPC1) transporter (*12*). These two membrane protein shares 23% identity and 44% similarity in protein sequence. Although the function of PfNCR1 is not completely clear, it has been proposed to be engaged in Chl trafficking (*11*). This gene is resistant to transposon mutagenesis (*13*) and a genetic knockdown study provides further evidence that *pfncr1* is an essential gene, critical for blood-stage parasite replication (*11*). Further, *pfncr1* knockdown causes the parasite to become hypersensitive to saponin, a pore-forming amphipathic glycoside with an affinity for free Chl. PfNCR1 localizes to the parasite plasma membrane at zones of contact with the parasitophorous vacuolar membrane that surrounds the parasite (*14*). Hence, inhibition via genetic knockdown of the PfNCR1 transporter leads to the accumulation of Chl on the parasite plasma membrane, indicating that PfNCR1 may be responsible for removing Chl from the parasite.

Taking together, PfNCR1 is an effective druggable target, as it is necessary for maintaining proper membrane lipid composition at the blood stage (*11*). To initiate our research efforts to uncover novel antimalarial drug targets and their action mechanisms to facilitate the development of antimalarial therapeutics, we decided to elucidate the structural basis of the PfNCR1 membrane protein. Our work detailed here provides molecular insights into the transport mechanism of Chl mediated by the PfNCR1 membrane protein to regulate membrane homeostasis. We present structures of PfNCR1 bound with Chl using single-particle cryo-electron microscopy (cryo-EM) to resolutions of 3.11 Å and 3.81 Å. We also report the 2.98 Å cryo-EM structure of PfNCR1 bound with MMV009108, a potential PfNCR1 inhibitor that heightens the sensitivity of *P. falciparum* to saponin. The structural information suggests that PfNCR1 contains two different binding sites for Chl and that this membrane protein forms a tunnel to shuttle of Chl between these two binding sites. It also enabled us to determine that MMV009108 creates a blockage within PfNCR1, forbidding it to shuttle Chl. Combined, cryo-EM, cholesterol efflux assays, molecular dynamics (MD) and target MD simulations, demonstrate that the PfNCR1 protein is capable of exporting cholesterols from the PPM to the parasitophorous vacuole (PV) to facilitate membrane homeostasis.

## Results

### Structural determination of the PfNCR1 transporter

To elucidate the structural information of PfNCR1, we cloned the gene *pfncr1* (PF3D7_0107500), which encodes the full-length *P. falciparum* PfNCR1 transporter of 1,470 amino acids, into the pcDNA3.1-N-DYK expression plasmid. PfNCR1 was expressed in HEK293 cells and purified using a strep-tactin affinity column. We further purified PfNCR1 using a superose 6 column and collected single-particle cryo-EM data of this membrane protein. Extensive classification of these single-particle images indicated that there were two distinct populations of PfNCR1 with different conformations coexisting in the single protein sample (Fig. S1). Several iterative rounds of classifications allowed us to sort the images based on these two distinct conformations. Three-dimensional reconstitutions of the two PfNCR1 transporter classes led to cryo-EM maps at nominal resolutions of 3.11 Å (PfNCR1-I) and 3.81 Å (PfNCR1-II) (Table S1 and Fig. S1), which enabled us to build two structural models of the PfNCR1 membrane protein to these resolutions. Each final structural model of the PfNCR1 transporter includes residues 3-180, 293-673, 719-736, 1046-1151 and 1169-1470.

### Structure of PfNCR1-I

Our cryo-EM structure indicates that PfNCR1 is monomeric in form with overall dimensions of 110 Å x 75 Å x 55 Å. This transporter consists of a large membrane-spanning domain formed by 12 transmembrane helices (TMs 1-12) and a large extension formed by two hydrophilic loops (loops 1 and 2), consistent with the signature of the resistance-nodulation-cell division (RND) superfamily of proteins (*15*) (Fig. 1A,B). It has been reported that the transmembrane domain of PfNCR1 resides at the parasite plasma membrane (PPM), while the two soluble loops protrude into and localize to the parasitophorous vacuole (PV) (*11*). These two large loops are located between TM1 and TM2, and between TM7 and TM8, where they contribute to subdomains PV1 and PV2 in the parasitophorous vacuolar space. Therefore, our cryo-EM structure indeed depicts that the PfNCR1 transporter contains two domains: the transmembrane PPM domain consisting of 12 TMs (TM1-12) and the soluble PV domain composed of subdomains (PV1 and PV2).

**Fig. 1.**
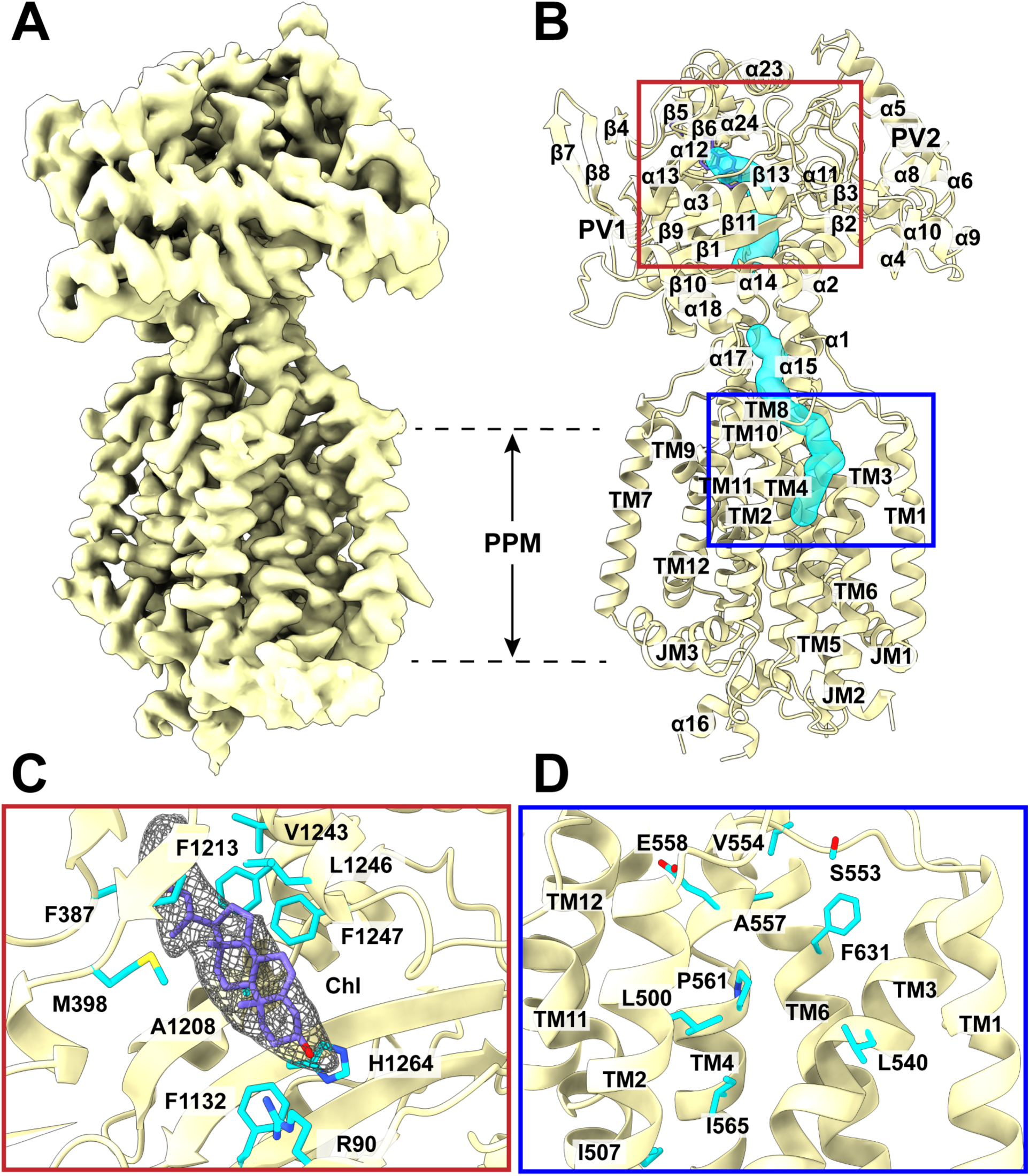
Structure of PfNCR1-I. (A) Cryo-EM map of PfNCR1-I at a resolution of 3.11 Å. (B) Side view of the ribbon diagram of PfNCR1-I viewed in the membrane plane. The structure of PfNCR1-I indicates that this transporter contains the PPM and PV domains. The PPM domain conisits of 12 TMs, whereas the PV domain is composed of subdomains PV1 and PV2. The PfNCR1 transporter forms a channel (cyan) spanning the PV domain and down to the outer leaflet of the PPM domain. This channel is broken into two halves. It appears residues 1105-1108, which form a short α17, block and split this tunnel into two portions. (C) The Chl binding site. Cryo-EM density of bound Chl is in gray meshes. The bound Chl molecule is in slate sticks. The binding residues R90, F387, M398, F1132, A1208, F1213, V1243, L1246, F1247 and H1264 are in cyan sticks. (D) The sterol sensitive domain (SSD). Residues L500, I507, L540, S553, V554, A557, E558, P561, I565 and F631 (cyan sticks) are found to surround the cavity formed at the SSD. These residues are expected to be important for substrate binding.

The N-terminal and C-terminal halves of PfNCR1 are assembled in a twofold pseudo-symmetrical fashion. These two halves can be superimposed, however, the root-mean-square-deviation (r.m.s.d.) of this superimposition is very high (15.9 Å for 386 Cα atoms), suggesting that the structures of these two halves are very different from each other. PV1 is composed of 16 α-helices and 10 β-strands (Fig. 1B). The majority of the PV1 amino acids emerge from loop 1. However, residues 1114-1125 of loop 2 also contribute to form helix α18 of PV1. PV2 constitutes nine α-helices and four β-strands (Fig. 1B). Similar to PV1, the PV2 amino acids mainly arise from loop 2, but residues 73-84, 142-161 and 295-298 of loop 1 participate in the formation of helices α2, α5 and α6 of this subdomain. The crossover of these two loops allows for the two subdomains to be spatially adjoined within the PV. Four flexible linkers, constituted by residues 85-88, 128-141, 300-335 and 1126-1130, are responsible for connecting these two subdomains together. Within the transmembrane domain of PfNCR1, two juxtamembrane helices (JM1 and JM2), approximately parallel to the PPM, are located at the N-terminus and between TM6 and TM7, respectively. These two JM helices may help strengthen the anchoring of this membrane protein in the PPM. The TMs are PPM embedded, but TM8 is significantly longer and protrude into the PV region. TM8 also directly tethers PV2 and forms part of the PV domain structure.

The TMs, JMs, α-helices and β-strands are designated numerically from the N-to C-termini: JM1 (4-34), TM1 (36-56), α1 (63-67), α2 (73-84), β1 (89-97), α3 (104-119), β2 (121-123), α4 (124-127), α5 (142-161), α6 (295-298), α7 (311-313), α8 (321-324), α9 (334-336), α10 (339-344), β3 (346-348), α11 (349-352), α12 (365-368), β4 (380-382), β5 (388-390), β6 (394-398), α13 (399-402), β7 (407-411), β8 (414-419), β9 (422-429), α14 (436-452), β10 (472-477), α15 (479-490), TM2 (494-514), TM3 (525-549), TM4 (555-582), TM5 (588-619), TM6 (623-659), JM2 (664-673), α16 (723-735), JM3 (1058-1073), TM7 (1075-1098), α17 (1105-1108), α18 (1114-1125), β11 (1131-1138), α19 (1140-1149), α20 (1185-1200), β12 (1204-1208), α21 (1210-1217), α22 (1221-1223), α23 (1227-1240), α24 (1242-1247), β13 (1260-1267), α25 (1273-1286), β14 (1297-1299), TM8 (1301-1334), TM9 (1338-1361), TM10 (1368-1394), TM11 (1400-1431) and TM12 (1435-1466).

A cleft is formed between subdomains PV1 and PV2. This cleft opens the top portion of the PV domain. A large cavity is also created at the gap between PV1 and PV2. The volume of this cavity is quite substantial and measured to be 6,156 Å^2^. Potentially, this cavity can form a substrate-binding site to accommodate PfNCR1 substrates. In addition to this PV domain cavity, PfNCR1 also possesses a large cavity at the PPM domain. This cavity is located at the outer leaflet of the lipid bilayer and surrounded by TMs 2, 3 and 4, which create an internal space of 345 Å^2^. In view of the structure, the architecture of these TMs resembles the sterol sensitive domain (SSD) found in both the human NPC1 (hNPC1) (*16, 17*) and PTCH1 (hPTCH1) (*18–20*) membrane proteins. Therefore, it is likely that that TMs 2-4 of PfNCR1 constitute a SSD cavity that is able to bind sterols and cholesterols.

Surprisingly, the structure indicates that PfNCR1 constitutes a tunnel-like feature starting from the outer leaflet of the PPM to the vacuolar cleft of the PV. This tunnel spans the two large internal cavities at the PPM and PV domains of the transporter (Fig. 1B). Potentially, this tunnel may allow for the transport of substrates across these two large cavities. However, this tunnel is broken into two halves, suggesting that the structure of PfNCR1-I may depict a conformational state with a closed form of this tunnel. The constriction site is created by two flexible loops from each side of the tunnel, where residues L482, Y1105, L1301 and Y1305 are responable for closing this tunnel (Fig. S2). Residues lining the wall of this tunnel, including F54, I63, L66, F92, M398, F477, L482, V486, I489, I492, L500, L501, V504, F536, F539, L540, P555, P556, A557, P561, F1109, F1132, A1208, L1246, F1247, L1301, F1305 and F1436, are found to be hydrophobic. Presumably, this tunnel could help facilitate the transport of hydrophobic substrates.

Unexpectedly, a large extra EM density was found in the structure of PfNCR1. This extra density is observed within the large space between PV1 and PV2, near the ceiling of the PV domain. The shape of this EM density resembles a large, elongated sterol molecule (Fig. 1C). To identify this fortuitous ligand, we employed gas chromatography coupled with mass spectrometry (GC-MS). GC-MS indicates that this ligand is cholest-5-en-3β-ol or cholesterol (Chl) (Fig. S3). Within 4.5 Å of the bound Chl molecule, there are at least 10 amino acids involved in the binding, R90, F387, M398, F1132, A1208, F1213, V1243, L1246, F1247 and H1264 (Fig. 1C). Notably, R90 forms a hydrogen bond with the hydroxyl group of this bound ligand. However, most of these residues that form the Chl binding site are hydrophobic in nature. Likewise, the cavity formed by the SSD of PfNCR1-I is surrounded by 10 residues, including L500, I507, L540, S553, V554, A557, E558, P561, I565 and F631, and many of them are hydrophobic resiudes (Fig. 1D).

Posttranslational oligosaccharide modifications have been found to play a critical role in localization, solubility and stability of many eukaryotic proteins. In the PfNCR1 structure, two extra cryo-EM densities are observed to directly connect to residues N165 and N294, indicating that these two aspargines may form N-linked glycosylation sites. Based on the cryo-EM densities, each asparagine is connected to a N-acetylglucosamine (NAG) moiety (Fig. S4). This structural feature is indeed in good agreement our liquid chromatography coupled with tandem mass spectrometry (LC-MS-MS) analysis, indicating that residues N165 and N294 are glycosylated (Fig. S5).

### Structure of PfNCR1-II

The overall conformation of this PfNCR1-II structure (Fig. 2A) is similar to that of PfNCR1-I. Superimposition of these two protomers results in an r.m.s.d. of 1.3 Å, indicating that these two PfNCR1 structures represent very different transient conformational states (Fig. 2B). A detailed inspection reveals that the elongated tunnel spanning the outer leaflet of the PPM domain and the PV domain is open (Fig. 2A and Fig. S2). Therefore, the conformation of PfNCR1-II most likely represents the open-tunnel form of this transporter. It appears the flexible linkers connecting the C-terminal end of TM7 and N-terminal end of α18 of PV1, and the C-terminal end of TM1 and N-terminal end of α2 of PV1 responsible for the opening and closing of this tunnel. In particular, residues 1105-1108 form a short α17 in PfNCR1-I and block this PfNCR1-I tunnel. In the PfNCR1-II structure, these residues switch its secondary structural conformation to form a random loop. In addition, the entire flexible linker formed by residues 1101-1110 of PfNCR1-II shifts its location and moves away from the central core of the tunnel by approximately 7 Å to open this PfNCR1-II tunnel (Fig. 2B).

**Fig. 2.**
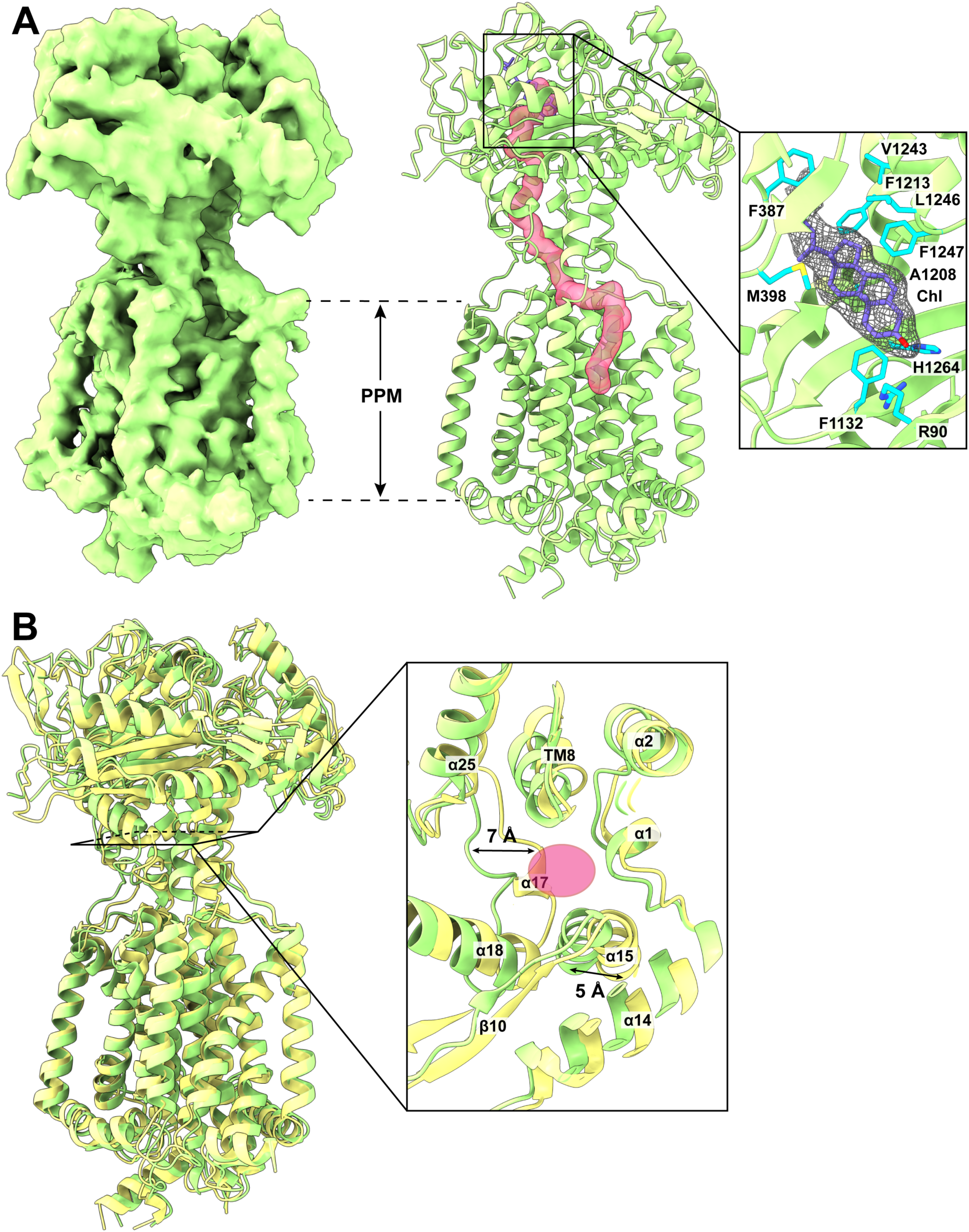
Structure of PfNCR1-II. (A) Cryo-EM map and side view of the ribbon diagram of PfNCR1-II viewed in the membrane plane at a resolution of 3.81 Å. Like PfNCR1-I, the structure of PfNCR1-II contains the PPM and PV domains. A continuous channel (pink) spanning the outer leaflet of the PPM domain and up to the PM domain. This tunnel is open in conformation, suggesting that the structure of PfNCR1-II represents the open-tunnel form of this transporter. A bound Chl molecule is found to occupy within the Chl-binding site between subdomains PV1 and PV2 of PfNCR1-II. This bound Chl is represented by slate sticks. The cryo-EM density of bound Chl is in gray meshes. Residues R90, F387, M398, F1132, A1208, F1213, V1243, L1246, F1247 and H1264, responsible for anchoring Chl are in cyan sticks. (B) Superimposition of the PfNCR1-I andPfNCR1-II structures. The secondary structural elements of PfNCR1-I are colored yellow, whereas those of PfNCR1-II are colored green. The cross sectional area of the narrowest region of the tunnel is also included. This cross section indicates that residues 1105-1108, which form a short α17 in PfNCR1-I (yellow), block and disconnect the tunnel of this transporter. In the PfNCR1-II structure (green), these residues switch its conformation to form a random coil and also shift away from the central core by approximately 7 Å to open the tunnel. The pink oval indicates the cross section of the narrowest region of the tunnel formed by PfNCR1-II.

Like PfNCR1-I, an extra density is found nearby the ceiling of the PV domain, corresponding to a bound Chl molecule sandwiched between subdomains PV1 and PV2 (Fig. 2A, insert). The location of bound Chl is more or less identical to that of PfNCR1-I. Within 4.5 Å of bound Chl, identical residues, including R90, F387, M398, F1132, A1208, F1213, V1243, L1246, F1247 and H1264, surround this sterol molecule to secure the binding.

### Structure of the PfNCR1-MMV009108 transporter-inhibitor complex

PfNCR1 has been shown to be a druggable target (*11*). It is required for maintaining the proper membrane lipid composition necessary for the development of parasites in the blood stage (*11*). A selection study utilizing isolated parasites with resistance-conferring mutations in PfNCR1 has allowed for the identification of three diverse antimalarial MMV small molecule compounds that directly interact with the PfNCR1 membrane protein. One such small molecule is MMV009108, which leads to the hypersensitivity of *P. falciparum* to saponins (*11*). To elucidate the structural basis of how PfNCR1 and MMV009108 interacts as well as the inhibition mechanism of this MMV compound, we incubated purified PfNCR1 with MMV009108 to form the PfNCR1-MMV009108 complex and then solved the cryo-EM structure of this complex to a resolution of 2.98 Å (Fig. 3A,B, Fig. S6 and Table S1).

**Fig. 3.**
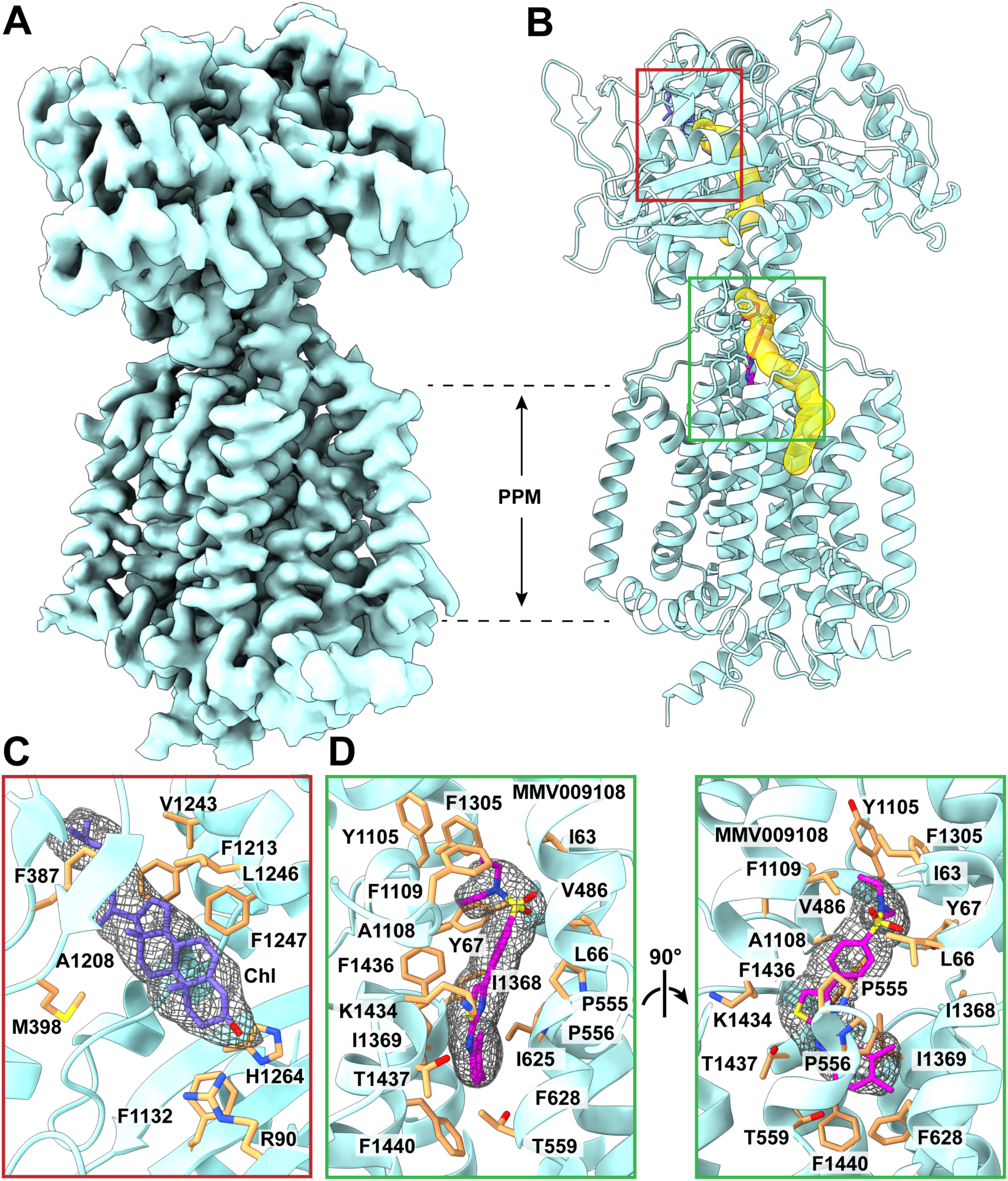
Structure of PfNCR1-MMV009108. (A) Cryo-EM map of PfNCR1-MMV009108 at a resolution of 2.98 Å. (B) Side view of the ribbon diagram of PfNCR1-MMV009108 viewed in the membrane plane. The structure of PfNCR1-MMV009108 indicates that this transporter forms a channel (yellow) spanning the outer leaflet of the PPM domain and up to the PV domain. Similar to PfNCR1-I, this channel is broken into two halves. (C) The Chl binding site. This Chl binding site is located between subdomains PV1 and PV2, identical to those found in the PfNCR1-I and PfNCR1-II structures. The bound Chl molecule is in slate sticks and its cryo-EM density is in gray meshaes. Residues R90, F387, M398, F1132, A1208, F1213, V1243, L1246, F1247 and H1264, which are important for Chl binding are in orange sticks. (D) The MMV009108 inhibitor binding site. The bound MMV009108 compound is in magenta sticks and its cryo-EM density is in gray meshes. Residues I63, L66, Y67, V486, P555, P556, T559, I625, F628, Y1105, A1108, F1109, F1305, I1368, I1369, K1434, F1436, T1437 and F1440, which are critical for interacting with MMV009108 are in orange sticks.

The overall structure of PfNCR1-MMV009108 is very similar to that of PfNCR1-I as described above. Superimposition of these two structures gives rise to a r.m.s.d. of 0.5 Å (for 871 Cα atoms), suggesting that the secondary structure of the transporter does not change significantly after inhibitor binding. Indeed, the structure of PfNCR1-MMV009108 depicts a closed tunnel conformation of the transporter, similar to that found in PfNCR1-I (Fig. S2). Interestingly, an extra density resembling Chl molecule is observed at the Chl-binding site (Fig. 3C). The mode of binding for this Chl is the same as that found in the PfNCR1 structure without the addition of inhibitor. Again, residues R90, F387, M398, F1132, A1208, F1213, V1243, L1246, F1247 and H1264 are responsible for the binding (Fig. 3C).

Our high quality cryo-EM map also allows us to unambiguously depict the location of bound MMV009108. This inhibitor binds within the tunnel created by the PfNCR1 transporter. Within 4.5 Å from bound MMV009108, residues I63, L66, Y67, V486, P555, P556, T559, I625, F628, Y1105, A1108, F1109, F1305, I1368, I1369, K1434, F1436, T1437 and F1440 are engaged in anchoring this inhibitor molecule (Fig. 3D). Based on this structural information, the inhibition mechanism of MMV009108 appears to be due to a physical blockage of the tunnel formed by PfNCR1, disallowing this transporter to export Chl from the PPM of the parasite. It is worth noting that residue A1108 is 3.6 Å away from the six-carbon aromatic ring of MMV009108, contacting this inhibitor via hydrophobic interaction. It has been documented that a mutation on this alanine to a threonine causes the MMV009108 drug to be incapable of inhibiting *P. falciparum*, reversing parasite susceptibility to saponins in the presence of this drug (*11*). Our structure of PfNCR1-MMV009108 is consistent with this inhibitor not binding the transporter in the A1108T mutant, possibly due to steric hinderance as well as electrostatic repulsion.

## Docking calculations

### Docking of Chl into PfNCR1

The cryo-EM structure of PfNCR1 depicts that this transporter creates two large cavities located at the vacuolar cleft of the PV and outer leaflet of the PPM, respectively. A bound Chl molecule was identified at the large cavity between subdomains PV1 and PV2. The second large cavity is surrounded by TMs 2, 3 and 4, mimicking the SSD-binding site at the outer leaflet of the PPM. We suspected that this SSD cavity is able to accommodate Chl. We, therefore, used the AutoDock Vina program (*21*) to elucidate if the SSD cavity located at the PPM is capable of binding Chl. We first docked Chl into the Chl-binding site formed between subdomains PV1 and PV2. We observed that Chl was bound at the same location as idenitifed in the cryo-EM structure of PfNCR1 (Fig. S7). We then studied the interaction of PfNCR1 with Chl at the SSD cavity using the same approach. We found that PfNCR1 specifically contacts and houses Chl in this SSD cavity surrounded by TMs 2-4, indicating that this cavity indeed forms a Chl-binding site (Fig. S7). The predicted binding affinities of Chl for these two Chl-binding sites are -6.9 and -8.5 kcal/mol, respectively.

### Docking of inhibitors into PfNCR1

We used the same approach to elucidate PfNCR1-inhibitor interactions based on AutoDock Vina (*21*). We first docked MMV009108 into its binding site that was found from cryo-EM. We found that PfNCR1 specific interacts with this inhibitor at the MMV009108 binding site as observed from the PfNCR1-MMV009108 structure (Fig. S8). We then studied the interactions of PfNCR1 with the other two inhibitors MMV028038 and MMV019662 using Vina (*21*). Vina indicated that both MMV028038 and MMV019662 specifically interact with PfNCR1 at their corresponding predicted binding sites along the cholesterol tunnel formed by PfNCR1 (Fig. S8). The predicted binding affinities for MMV009108, MMV028038 and MMV019662 with PfNCR1 were calculated to be -13.2, -10.1 and -12.2 kcal/mol, respectively.

### Computational simulations of the PfNCR1 transporter

The gene *pfncr1* (PF3D7_0107500), encoding the PfNCR1 membrane protein, is annotated as a lipid/sterol transporter. Based on the structural information, PfNCR1 creates two Chl-binding sites and an elongated tunnel spanning these two sites. It is likely that this transporter is capable of shuttling Chl from one binding sites to the other. To elucidate the mechanism of Chl transport, we performed molecular dynamics (MD) (Fig. S9 and Fig. S10) and target MD (Fig. S11) simulations on the PfNCR1 transporter in an explicit lipid bilayer and water environment using Amber (*22, 23*) and NAMD (*24*).

### Dynamics of the PfNCR1 transporter

Principle component analysis (PCA) indicates that the first and second eigenvectors, which depict the two most important motions extracted from the MD simulation trajectory, correspond to a rigid-body movement of subdomain PV1 in relation to subdomain PV2 of the PfNCR1 transporter (Fig. S10). This rigid-body motion is accompanied by slightly movements of the TM helices at the PPM domain. Interestingly, the rigid-body movement of the PV1 subdomain can be interpreted as the motion to open and close the cleft between subdomains PV1 and PV2. This motion could help promote the transport of substrates via the PfNCR1 transporter.

### Cholesterol transport pathway

Target MD simulations allowed us to observe that the Chl molecule is able to follow the path of the tunnel identified from the cryo-EM structure of PfNCR1 and shuttle from the PPM to the PV region (Fig. S11). According to the simulations, Chl first enters the PfNCR1 transporter via the Chl-binding site at the SSD cavity surrounded by TMs 2-4. It then passes through the elongated tunnel connecting the PPM and PV regions, and then reaches PV cleft to the Chl-binding site located between subdomains PV1 and PV2. Target MD simulations indicate that PfNCR1 is capable of exporting Chl from the outer leaflet of the PPM to the PV region.

### Putative proton transfer pathway

We performed 1 μs MD simulations on PfNCR1-I. We identified two water-accessible pockets, as illustrated in Fig. S12. The first pocket, situated at the outer leaflet side of the PPM, is composed of residues D1352, S1370, S1377, S1381, T1443 and S1451. The second pocket, located at the inner leaflet side of the PPM, is formed by residues Y510, D570, D571, D1383, H1384 and H1387. These water-accessible residues likely play a crucial role in forming the proton-relay network for energy coupling. Fig. S12 illustrates the detailed putative proton transfer pathway. Based on the simulation, protons enter the outer leaflet side of the water-accessible pocket and subsequently transfer through the proton-relay residues D1352, S1370, S1377, S1381, T1443 and S1451. D1383, H1384, S1451, and S1381. The pathway also involves passing through residues D1383, H1384 and H1387, enabling entry into the inner leaflet side of the water-accessible pocket, and then to residues D570, D571 and Y510, where these protons finally reach the cytoplasmic region of the parasite.

### Cholesterol efflux assays

HEK293 cells were used to generate an inducible PfNCR1 stable cell line in the presence of tetracycline (+ tet) (Fig. S13). This stable line was exposed to fluorescently-labeled cholesterols (NBD-cholesterol, 22-(*N*-(7-nitrobenz-2-oxa-1,3-diazol-4-yl)amino)-23,24-bisnor-5-cholen-3β-ol) in order to load the NBD-cholesterol molecules onto the cell membrane. We then individually conducted comparative analyses of NBD-cholesterol efflux in the control (-tet) and PfNCR1-expressing HEK293 cells (+ tet) at three different extracellular pHs (pH = 7.0, pH = 7.4 and pH = 8.0). At the extracellular pH of 7.0, a pH lower than that of the intracellular pH, pH = 7.4, the efflux of NBD-cholesterol from the plasma membrane was markedly increased in cells expressing PfNCR1 (+ tet) when compared with that of the control HEK293 cells (-tet) as indicated by the rapid decrease in fluorescence signal (Fig. 4A). The results also indicated that cells expressing PfNCR1 generated an exponential fluorescence decay curve with the rate of 0.39 ± 0.04 s^-1^. This fluorescence decay rate is much faster than that of the control cells (0.10 ± 0.02 s^-1^), suggesting that the expression of PfNCR1 facilitates the export of NBD-cholesterol from the membrane. However, when the extracellular pH was adjusted to either 7.4 or 8.0, no difference in the efflux of NBD-cholesterol from the plasma membrane was found, as the fluorescence curves of HEK293 cells with and without expressed PfNCR1 overlap with each other (Fig. 4B, C). These experimental data suggest that the expression of PfNCR1 facilitates the efflux of cholesterols from the cell membrane, and that this process is pH dependent where the influx of protons may be a prerequisite to provide the proton-motive-force (PMF) needed for cholesterol removal.

**Fig. 4.**
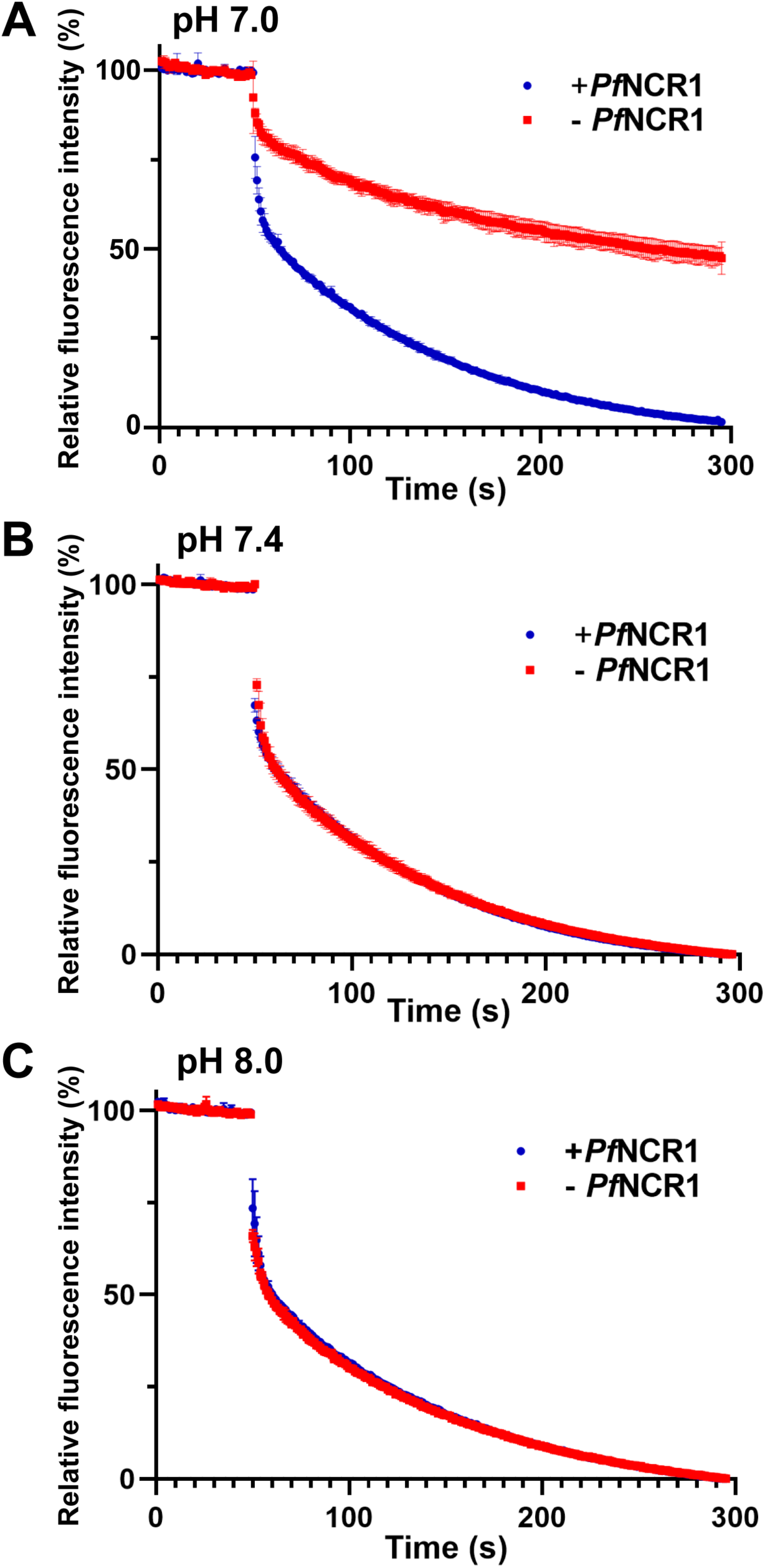
PfNCR1 facilitates the efflux of cholesterol from mammalian cells. (A) At the extracellular pH of 7.0, the efflux of NBD-cholesterol from the plasma membrane of HEK293 cells expressing PfNCR1 (+ tet) is markedly increased as indicated by the rapid decrease in fluorescence signal when compared with that of the control HEK293 cells (-tet). (B) At the extracellular pH of 7.4, the efflux of NBD-cholesterol from the plasma membrane of HEK293 cells expressing PfNCR1 (+ tet) was found to be no different from that of the control HEK293 cells (-tet), where the fluorescence curves of HEK293 cells with and without expressed PfNCR1 overlap with each other. (C) At the extracellular pH of 8.0, the efflux of NBD-cholesterol from the plasma membrane of HEK293 cells expressing PfNCR1 (+ tet) was found to be no different from that of the control HEK293 cells (-tet), similar to that found in (B). In (A-C), all experiments were repeated by three times (blue, HEK293 cells expressing PfNCR1; red, control HEK293 cells withour expressing PfNCR1). The small error bars depict standard deviation for *n* = 3. In (A), all fluorescence decay data points from HEK293 cells expressing PfNCR1 at the time interval between 5 and 30 minutes are significantly different from that of control cells without expressing PfNCR1 (*P* < 0.0001; student’s *t*-test).

## Discussion

In this study, we successfully solved high-resolution cryo-EM structures of the PfNCR1 transporter both in the absence and presence of the inhibitor MMV009108. This transporter has been proven to be an attractive antimalarial target, as it resides at the parasite’s PPM and plays an important functional role during intraerythrocytic growth. Knockdown of the *pfncr1* gene or drug inhibition of the PfNCR1 membrane protein is lethal to the parasites and increases susceptibility of the PPM to cholesterol-intercalating saponin glycosides. PfNCR1 in our structure is seen to bind cholesterol, which helps explain the observed phenotypic effects. The saponin sensitivity likely reflects aberrant cholesterol accumulation in the PPM when PfNCR1 is impaired. Digestive vacuoles that form from the parasite membranes are normally devoid of cholesterol (*11, 25*) but in the knockdown/inhibited parasites form aberrant structures with abnormal membrane curvature, again suggesting cholesterol accumulation. It seems likely that the role of PfNCR1 is to remove Chl from the PPM. Our cryo-EM structure of PfNCR1 indicates that a bound Chl molecule is found within the cavity created between subdomains PV1 and PV2. Moreover, an elongated tunnel within the transporter directly connects this Chl-binding site to the SSD cavity. Autodock Vina suggests that the SSD cavity is capable of binding Chl, creating a putative Chl-binding site. Cholesterol efflux assays indicate that recombinant PfNCR1 expressed in HEK293 cells is able to facilitate the export and removal of Chl from the cell membrane. This observation is further strengthened by target MD simulations, where PfNCR1 is capable of transporting Chl from the PPM to PV domains of PfNCR1 utilizing the tunnel formed within this transporter.

Based on the structural, computational and biochemical results, we propose that the PfNCR1 transporter captures Chl from the outerleaflet of the PPM, shuttles it to the SSD and PV Chl-binding sites, and eventually exports the sterol molecule to the parasitophorous vacuolar space via the Chl-tunnel constituted by the transporter (Fig. 5). A cleft surrounded by subdomains PV1 and PV2 of the PV domain forms the Chl-binding site. This binding site located at the PV domain probably also marks the exit site of the tunnel, which spans the SSD cavity situated in the outer leaflet of the PPM and up to the parasitophorous vacuolar space. In the parasite, the PPM and PVM form regions of close contact, which would allow for lipid transfer (*14*). PfNCR1 localizes to these sites while proteins involved in aqueous solute transport localize to separate domains in the PVM. As a member of the resistance-nodulation-cell division (RND) superfamily of transporters (*15*), PfNCR1 is likely a proton-motive-force (PMF)-dependent transporter that functions via a substrate/proton antiport mechanism. Therefore, PfNCR1 may get its energy to work against the concentration gradient and remove Chl from the Chl-poor PPM to the Chl-rich PVM using the PMF. This would require the protons being relayed in the opposite direction of Chl efflux. Two independent studies comparing parasite and infected erythrocyte cytoplasmic pH have been done, with similar results (*26, 27*). The pH external to the parasite was estimated at 7.1 (one measured 6.9 close to the parasite) and the pH inside the parasite was 7.3. Thus, a gradient of 0.2 to 0.4 pH units exists across the parasite surface so the directionality of proton transfer should be the influx of protons from the erythrocyte and contiguous parasitophorous vacuolar space to the cytoplasm of the parasite. Our MD simulations data indeed suggest a putative proton-relay network that facilitates the transport of protons in this direction. Under these pH conditions, our cholesterol efflux assays indicate that PfNCR1 is able to mediate the efflux of Chl from the parasite’s plasma membrane to the parasitophorous vacuolar space. This would be similar to what is seen in the bacterial RND superfamily of transport systems, including the hopanoid lipid transporter *Burkholderia multivorans* HpnN (*28*) and the trehalose monomycolate lipid transporter *Mycobacterium smegmatis* MmpL3 (*29, 30*), where these transporters are responsible for bacterial cell envelope and cell membrane biogenesis.

**Fig. 5.**
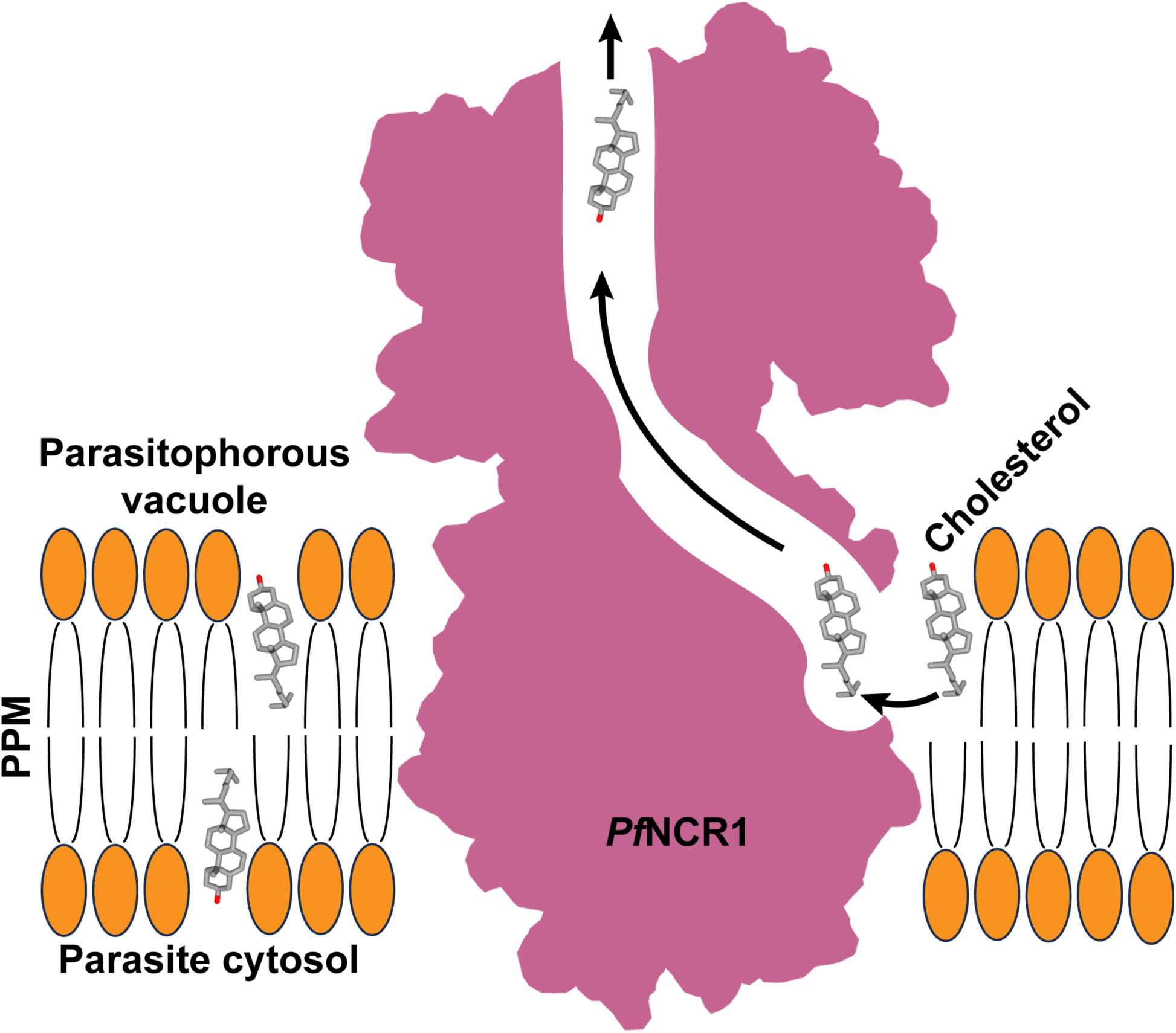
Proposed mechanism for Chl translocation via PfNCR1. This schematic diagram indicates that the PfNCR1 transporter is capable of picking up a Chl molecule from the parasite’s plasma membrane. This Chl molecule will arrive the SSD of PfNCR1 and pass through the tunnel formed by this transporter. The Chl moiety will then reach the cholesterol binding site located between PV1 and PV2 of the PV domain of PfNCR1 in the parasitophorous vacuole and be exported out of the transporter.

PfNCR1 is an attractive novel antimalarial drug target. Its druggability has been demonstrated by the inhibitory efficacy of three MMV drug candidates. When these compounds directly interact with PfNCR1, they elevate the sensitivity of *P. falciparum* to saponin lysis. Our data indeed demonstrate that these drugs specifically bind to functional motifs within the transporter and that their inhibition mechanism is via a direct blockage of the PfNCR1 tunnel, hindering the function of this transporter to traffic Chl. As the expression of PfNCR1 is capable of mediating the parasite’s resistance to saponin, it raises the question whether PfNCR1 can also function as a drug efflux pump specific to steroidal antimalarials. A similar phenomenon has been seen in the AbgT family of bacterial membrane proteins, where these transporters have been found to be capable of exporting the folate synthesis catabolite *p*-aminobenzoyl-glutamate (PABA) from bacterial cells (*31–33*). However, their canonical function is to perform as antibiotic efflux pumps and extrude sulfonamides to mediate bacterial resistance to these antimetabolite drugs (*31–33*).

Our study is a significant advancement in the field of antimalarial therapeutics as it defines vulnerable functional properties in the structure of PfNCR1 in the context of its herein identified substrate Chl. These findings can be used to develop novel antimalarial drugs that are unlikely to be sensitive to current parasite drug resistance mechanisms.

## Materials and Methods

### Expression and purification of PfNCR1

The codon-optimized DNA of full-length PfNCR1 from *Plasmodium falciparum* PF3D7_0107500 was synthesized and cloned into pcDNA3.1-N-DYK (GenScript) in frame with a thrombin cleavage site and Twin-Strep-tag at the C-terminus. The resulting plasmid was confirmed by the Sanger method of DNA sequencing.

The human embryonic kidney Expi293F (Thermo Fisher) cells were cultured in Expi 293^TM^ Expression Medium (Gibco) at 37°C supplemented with 8% CO_2_. The PfNCR1 protein was expressed using a transient expression system with the following procedures. The purified pcDNA3.1-N-DYK plasmid expressing full-length PfNCR1 was mixed with cationic liposomes (Transfection reagent I, Avanti Polar lipid) at a 1:10 (DNA:liposome) (w/w) ratio in Opti-MEM™ I reduced-serum medium (Gibco) and incubated at room temperature for 15 min. The resulting lipoplexes were added to cells (cell density of 2.5-3×10^6^ cells/ml) at a final DNA concentration of 1 mg/l. To boost protein expression, valproic acid (Sigma) was added at a final concentration of 3 mM after 18-24 h and allowed to proceed for total 72 h.

Cells were collected and resuspended in lysis buffer (20 mM HEPES-NaOH (pH 7.5) and 150 mM NaCl) and disrupted with a French pressure cell. The membrane fraction was collected and washed once with the lysis buffer. The membrane protein was then solubilized in 1% (w/v) glycol-diosgenin (GDN) for 3 h at 4°C. Insoluble material was removed by ultracentrifugation at 100,000 x g. The extracted protein was applied to a Strep-Tactin affinity column (IBA Lifesciences) and washed twice with 15 column volumes of lysis buffer supplemented with 0.01% GDN. The PfNCR1 protein was eluted by adding 4 mM desthiobiotin. The purity of the PfNCR1 protein (>90%) was judged using SDS-PAGE stained with Coomassie Brilliant Blue. To enhance sample homogeneity, the PfNCR1 protein was further purified using a superose 6 column (GE Healthcare) equilibrated with 20 mM Tris-HCl, pH 7.5, 100 mM NaCl and 0.005% GDN. The purified protein was then concentrated to a final concentration of 7 mg/ml.

### Electron microscopy sample preparation

The PfNCR1 protein embedded in GDN detergent micelles was concentrated to 7 mg/ml. A 2.5 μl sample was applied to glow-discharged holey carbon grids (Quantifoil Cu R1.2/1.3, 300 mesh), blotted for 5 s and then plunge-frozen in liquid ethane using a Vitrobot (Thermo Fisher). The grids were transferred into cartridges. For high resolution data collection, the sample grids were loaded into a Titan Krios cryo-electron microscope operated at 300 kV equipped with Gatan BioQuantum imaging filter (GIF) and a K3 summit direct electron detector (Gatan). The micrographs were recorded using SerialEM (*34*) with counting mode at nominal x 81,000 magnification corresponding to a calibrated pixel size of 1.07 Å (super-resolution, 0.535 Å/pixel) and a defocus range of -1 to -1.5 μm. To remove inelastically scattered electrons, the slit width was set to 20 eV. Each micrograph was exposed with a total specimen dose between 33 aand 35.9 e^-^/Å^2^ and ∼38 frames were captured per specimen area.

For the PfNCR1-MMV009108 inhibitor complex, a 40 μM PfNCR1 sample was incubated with 50 μM MMV009108 for 2 h to form the PfNCR1-MMV009108 complex. A 1 μl sample was applied to glow-discharged holey carbon grids (UltrAufoil Au R1.2/1.3, 300 mesh), blotted for 5 s, and a 1.7 μl sample was applied to the grid again with 3 s blotting, and then plunge-frozen in liquid ethane using a Vitrobot (Thermo Fisher). The procedures for high-resolution cryo-EM data collection were the same as those described above. Each micrograph was exposed with a total specimen dose of 40.5 e^-^/Å^2^ and 45 frames were captured per specimen area.

### Cryo-EM data processing

The micrographs of PfNCR1 were aligned by using patch-based motion correction for beam induced motion using CryoSPARC (*35*). The contrast transfer function (CTF) parameters of the micrographs were determined using Patch CTF (*36*). After manual inspection and sorting to discard poor micrographs, ∼2,000 particles of PfNCR1 were manually picked to generate templates for automatic picking. Initially, 6,977,953 particles were selected after autopicking in CryoSPARC. Several iterative rounds of 2D classifications followed by ab initio and heterogeneous 3D classifications were performed to remove false picks and classes with unclear features, ice contamination or carbon. The 3D classification analysis was then employed, resulting in two distinct classes. Non-uniform refinement followed by local refinement with non-uniform sampling resulted in 3.11 Å- and 3.81 Å-resolution cryo-EM maps for PfNCR1-I and PfNCR1-II based on gold standard Fourier shell correlation (FSC 0.143).

### Model building and refinement

Model building of PfNCR1-I was based on the 3.11 Å cryo-EM map. The predicted modeling structure of PfNCR1 generated by AlphaFold (*37*) was fitted into the density map using Chimera (*38*). Subsequent model rebuilding was performed using Coot (*39*). Structure refinements were performed using the phenix.real_space_refine program (*40*) from the PHENIX suite (*41*). The final atomic model was evaluated using MolProbity (*42*). Statistics associated with data collection, 3D reconstruction, and model refinement are included in Table S1.

The PfNCR1-MMV009108 structural model at 2.98 Å resolution was built based on the PfNCR1-I structure. A 3D conformer of the inhibitor MMV009108 (Pub Chem CID: 2297659) was processed using phenix.elbow implemented in PHENIX (*41*). Structural refinements were done using the same approach as described above (Table S1).

### Molecular docking

The program AutoDock Vina (*21*) was used to predicted the binding modes of Chl and the three PfNCR1 inhibitors, including MMV009108, MMV028038 and MMV019662. The PfNCR1-I structure (with the Chl molecule removed) was used for dockings. The protein was set as a rigid structure, whereas the conformation of each antibiotic molecule was optimized via all modeling and docking procedures. For each inhibitor, the results were ranked on the basis of predicted free binding energy, where the one with the highest binding affinity was recorded.

### Molecular dynamics (MD) simulations

The protonation states of the titratable residues of the PfNCR1 transporter was determined using the H++ server (http://biophysics.cs.vt.edu/). The cryo-EM structure of PfNCR1-I was used as the template for the PfNCR1-Chl structure. We also removed the bound Chl ligand from the PfNCR1-I structure to form the template for apo-PfNCR1. These two structures were individually immersed in an explicit lipid bilayer consisting of POPC, POPE, POPS and Chl with a molecular ratio of 25:5:5:1 using the CHARMM-GUI Membrane Builder webserver (http://www.charmm-gui.org/?doc=input/membrane). A water box with dimensions of 104.9 Å x 106.7 Å x 170.0 Å was employed. 150 mM NaCl and extra neutralizing counter ions were added for these simulations. The total number of atoms were 144,403 and 144,432 for the apo-PfNCR1 and PfNCR1-Chl systems, respectively. The Antechamber module of AmberTools was employed to generate parameters for Chl using the general AMBER force field (GAFF) (*22, 23*). The partial charges of Chl were calculated using *ab initio* quantum chemistry at the HF/6-31G* level (GAUSSIAN 16 program) (Gaussian Inc., Wallingford). The RESP charge-fitting scheme was used to calculate partial charges on the atoms (*43*). The tleap program was used to generate parameter and coordinate files using the ff14SB and Lipid17 force field for both the protein and lipids. The PMEMD.CUDA program implemented in AMBER18 (AMBER 2018, UCSF) was used to conduct MD simulations. The simulations were performed with periodic boundary conditions to produce isothermal-isobaric ensembles. Long range electrostatics were calculated using the particle mesh Ewald (PME) method (*44*) with a 10 Å cutoff. Prior to production runs, energy minimization of these systems was carried out. Subsequently, the systems were heated from 0 K to 303 K using Langevin dynamics with 1 ps^-1^ collision frequency. During heating, the PfNCR1 transporter was position-restrained using an initial constant force of 500 kcal/mol/Å^2^ and weakened to 10 kcal/mol/Å^2^ to allow for the movement of lipid and water molecules. Then, the systems went through 5 ns equilibrium MD simulations. Finally, a total of 1 ms production MD simulations were conducted. During simulations, the coordinates were saved in every 500 ps for analysis. All systems were well equilibrated after 100 ns simulations according to root mean square deviations (RMSD) of the transporter’s Cα atoms. 100 ns – 1 ms trajectories of each system were used for root mean square fluctuation (RMSF) and principal component analysis (PCA) (*45, 46*). GROMCAS analysis tools were used for the MD simulation trajectory analysis (*47*).

### Target MD simulations

Target MD (TMD) was performed using the NAMD program (*24*) with the same AMBER force field parameters as described above. In the simulations, we selected the heavy atoms of Chl to be guided towards the target position (from the PPM domain to the PV domain) by the application of steering forces. The root mean square (RMS) distance between the current coordinates and the target structure was calculated at each timestep. The force on each selected atom was given by a gradient of potential as a function of the RMS values. The system was gone through energy minimization, heating, and 5 ns equilibrium MD simulations. Then, TMD simulation was performed for 5 ns based on the MD equilibrated coordinates. A value of 500 kcal/mol/Å^2^ was used as an elastic constant for TMD forces during the simulations.

### Generation of PfNCR-1-expressing stable cell line

PfNCR1 containing a C-terminal Strep epitope was stably expressed in the HEK293 Flp-In T-REx cell line following the manufacturer’s protocol (Invitrogen). Cells were cultured in DMEM medium supplemented with 5% FBS and 1% PenStrep (Invitrogen). Tetracycline induction at a concentration of 1000 ng/ml was applied for 40 hours to prompt the expression of PfNCR1, as confirmed by western blot analysis. Controls without tetracycline induction were incorporated into each experiment.

### NBD-cholesterol export

PfNCR1-expressing HEK293 Flp-In T-REx cells were induced for 40 h and then starved overnight (16 h) to deplete endogenous cholesterol. This was followed by a 2 h incubation in 2 mM MCD and 5 μM NBD-cholesterol. Subsequently, cells were removed from the plate and washed three times with *N*-2-hydroxyethylpiperazine-*N*′-2-ethanesulfonic acid (HEPES)-Tyrode’s buffer (pH 7.4) (10 mM HEPES, 12 mM NaHCO_3_, 130 mM NaCl, 5 mM D-glucose, 5 mM KCl, 0.4 mM NaHPO_4_, 1 mM MgCl_2_) and diluted to 1 × 10^6^ cells per ml.

NBD-cholesterol efflux was performed using a PTI QuantaMaster 4/2005 fluorometer (Photon Technology International, Inc.) at 20 °C with 1 s each interval for 300 s. After the first 50 s, 10 mM dithionite was added to the cell suspension, which was continuously stirred using a small magnetic stirrer with dimensions of 6 mm x 1.5 mm x 1.5 mm, to quench the fluorescence signal of unprotected NBD-cholesterols in solution. The excitation and emission wavelengths were 469 nm and 537 nm, respectively. The experiments were repeated three times, including those with control cells and cells expressing PfNCR1.

## Data availability

Atomic coordinates and structure factors have been deposited with accession codes 8V0G (PDB) and EMD-42854 (EMBD) for PfNCR1-I; 8V12 (PDB) and EMD-42878 (EMDB) for PfNCR1-II; 8V1G (PDB) and EMD-42885 (EMDB) for PfNCR1-MMV009108.

## References

1. WHO. (World Health Organization, Geneva, Switzerland, 2022).

2. CDC. (Centers for Disease Control and Prevention, Atlanta, GA, USA, 2018).

3. B. Blasco, D. Leroy, D. A. Fidock, Antimalarial drug resistance: linking Plasmodium falciparum parasite biology to the clinic. Nat Med 23, 917–928 (2017).

4. S. Lauer et al., Vacuolar uptake of host components, and a role for cholesterol and sphingomyelin in malarial infection. Embo J 19, 3556–3564 (2000).

5. A. G. Maier, C. van Ooij, The role of cholesterol in invasion and growth of malaria parasites. Front Cell Infect Microbiol 12, 984049 (2022).

6. M. Koch et al., The effects of dyslipidaemia and cholesterol modulation on erythrocyte susceptibility to malaria parasite infection. Malar J 18, 381 (2019).

7. N. D. Geoghegan et al., 4D analysis of malaria parasite invasion offers insights into erythrocyte membrane remodeling and parasitophorous vacuole formation. Nat Commun 12, 3620 (2021).

8. A. I. Ahiya, S. Bhatnagar, J. M. Morrisey, J. R. Beck, A. B. Vaidya, Dramatic Consequences of Reducing Erythrocyte Membrane Cholesterol on Plasmodium falciparum. Microbiol Spectr 10, e0015822 (2022).

9. A. R. Dluzewski, K. Rangachari, R. J. Wilson, W. B. Gratzer, Relation of red cell membrane properties to invasion by Plasmodium falciparum. Parasitology 91, 273–280 (1985).

10. S. Bhatnagar, S. Nicklas, J. M. Morrisey, D. E. Goldberg, A. B. Vaidya, Diverse Chemical Compounds Target Plasmodium falciparum Plasma Membrane Lipid Homeostasis. ACS Infect Dis 5, 550–558 (2019).

11. E. S. Istvan et al., Plasmodium Niemann-Pick type C1-related protein is a druggable target required for parasite membrane homeostasis. Elife 8, e40529 (2019).

12. P. G. Pentchev, Niemann-Pick C research from mouse to gene. Biochim Biophys Acta 1685, 3–7 (2004).

13. M. Zhang et al., Uncovering the essential genes of the human malaria parasite Plasmodium falciparum by saturation mutagenesis. Science 360, eaap7847 (2018).

14. M. Garten et al., Contacting domains segregate a lipid transporter from a solute transporter in the malarial host-parasite interface. Nat Commun 11, 3825 (2020).

15. T. T. Tseng et al., The RND permease superfamily: an ancient, ubiquitous and diverse family that includes human disease and development proteins. J Mol Microbiol Biotechnol 1, 107–125 (1999).

16. X. Li et al., Structure of human Niemann-Pick C1 protein. Proc Natl Acad Sci U S A 113, 8212–8217 (2016).

17. X. Gong et al., Structural Insights into the Niemann-Pick C1 (NPC1)-Mediated Cholesterol Transfer and Ebola Infection. Cell 165, 1467–1478 (2016).

18. X. Gong et al., Structural basis for the recognition of Sonic Hedgehog by human Patched1. Science 361, eaas8935 (2018).

19. X. Qi, P. Schmiege, E. Coutavas, J. Wang, X. Li, Structures of human Patched and its complex with native palmitoylated sonic hedgehog. Nature 560, 128–132 (2018).

20. Y. Zhang et al., Structural Basis for Cholesterol Transport-like Activity of the Hedgehog Receptor Patched. Cell 175, 1352–1364 (2018).

21. O. Trott, A. J. Olson, AutoDock Vina: improving the speed and accuracy of docking with a new scoring function, efficient optimization, and multithreading. J Comput Chem 31, 455–461 (2010).

22. J. M. Wang, R. M. Wolf, J. W. Caldwell, P. A. Kollman, D. A. Case, Development and testing of a general amber force field. Journal of Computational Chemistry 25, 1157–1174 (2004).

23. J. M. Wang, W. Wang, P. A. Kollman, D. A. Case, Automatic atom type and bond type perception in molecular mechanical calculations. Journal of Molecular Graphics & Modelling 25, 247–260 (2006).

24. J. C. Phillips et al., Scalable molecular dynamics with NAMD. Journal of Computational Chemistry 26, 1781–1802 (2005).

25. J. M. Matz, Plasmodium’s bottomless pit: properties and functions of the malaria parasite’s digestive vacuole. Trends Parasitol 38, 525–543 (2022).

26. S. Wünsch et al., Differential stimulation of the Na+/H+ exchanger determines chloroquine uptake in Plasmodium falciparum. J Cell Biol 140, 335–345 (1998).

27. M. Hayashi et al., Vacuolar H(+)-ATPase localized in plasma membranes of malaria parasite cells, Plasmodium falciparum, is involved in regional acidification of parasitized erythrocytes. J Biol Chem 275, 34353–34358 (2000).

28. N. Kumar et al., Crystal structures of the Burkholderia multivorans hopanoid transporter HpnN. Proc Natl Acad Sci U S A 114, 6557–6562 (2017).

29. C. C. Su et al., MmpL3 is a lipid transporter that binds trehalose monomycolate and phosphatidylethanolamine. Proc Natl Acad Sci U S A 116, 11241–11246 (2019).

30. C. C. Su et al., Structures of the mycobacterial membrane protein MmpL3 reveal its mechanism of lipid transport. PLoS Biol 19, e3001370 (2021).

31. J. R. Bolla et al., Crystal structure of the Alcanivorax borkumensis YdaH transporter reveals an unusual topology. Nat Commun 6, 6874 (2015).

32. C. C. Su et al., Structure and function of Neisseria gonorrhoeae MtrF illuminates a class of antimetabolite efflux pumps. Cell Rep 11, 61–70 (2015).

33. J. A. Delmar, E. W. Yu, The AbgT family: A novel class of antimetabolite transporters. Protein Sci 25, 322–337 (2016).

34. D. N. Mastronarde, Automated electron microscope tomography using robust prediction of specimen movements. J Struct Biol 152, 36–51 (2005).

35. A. Punjani, J. L. Rubinstein, D. J. Fleet, M. A. Brubaker, cryoSPARC: algorithms for rapid unsupervised cryo-EM structure determination. Nat Methods 14, 290–296 (2017).

36. K. Zhang, Gctf: Real-time CTF determination and correction. J Struct Biol 193, 1–12 (2016).

37. J. Jumper et al., Highly accurate protein structure prediction with AlphaFold. Nature 596, 583–589 (2021).

38. E. F. Pettersen et al., UCSF Chimera--a visualization system for exploratory research and analysis. J Comput Chem 25, 1605–1612 (2004).

39. P. Emsley, K. Cowtan, Coot: model-building tools for molecular graphics. Acta Crystallogr D Biol Crystallogr 60, 2126–2132 (2004).

40. P. V. Afonine et al., Real-space refinement in PHENIX for cryo-EM and crystallography. Acta Crystallogr D Struct Biol 74, 531–544 (2018).

41. P. D. Adams et al., PHENIX: building new software for automated crystallographic structure determination. Acta Crystallogr D Biol Crystallogr 58, 1948–1954 (2002).

42. V. B. Chen et al., MolProbity: all-atom structure validation for macromolecular crystallography. Acta Crystallogr D Biol Crystallogr 66, 12–21 (2010).

43. C. M. Breneman, K. B. Wiberg, DETERMINING ATOM-CENTERED MONOPOLES FROM MOLECULAR ELECTROSTATIC POTENTIALS - THE NEED FOR HIGH SAMPLING DENSITY IN FORMAMIDE CONFORMATIONAL-ANALYSIS. Journal of Computational Chemistry 11, 361–373 (1990).

44. T. Darden, D. York, L. Pedersen, PARTICLE MESH EWALD - AN N.LOG(N) METHOD FOR EWALD SUMS IN LARGE SYSTEMS. Journal of Chemical Physics 98, 10089–10092 (1993).

45. A. Amadei, A. B. Linssen, H. J. Berendsen, Essential dynamics of proteins. Proteins 17, 412–425 (1993).

46. A. Amadei, M. A. Ceruso, A. Di Nola, On the convergence of the conformational coordinates basis set obtained by the essential dynamics analysis of proteins’ molecular dynamics simulations. Proteins-Structure Function and Genetics 36, 419–424 (1999).

47. H. J. C. Berendsen, D. Vanderspoel, R. Vandrunen, GROMACS - A MESSAGE-PASSING PARALLEL MOLECULAR-DYNAMICS IMPLEMENTATION. Computer Physics Communications 91, 43–56 (1995).

