## Supplementary Figures and Tables for "The Plasmodium falciparum NCR1 membrane protein is a novel antimalarial target that exports cholesterol to maintain membrane homeostasis"

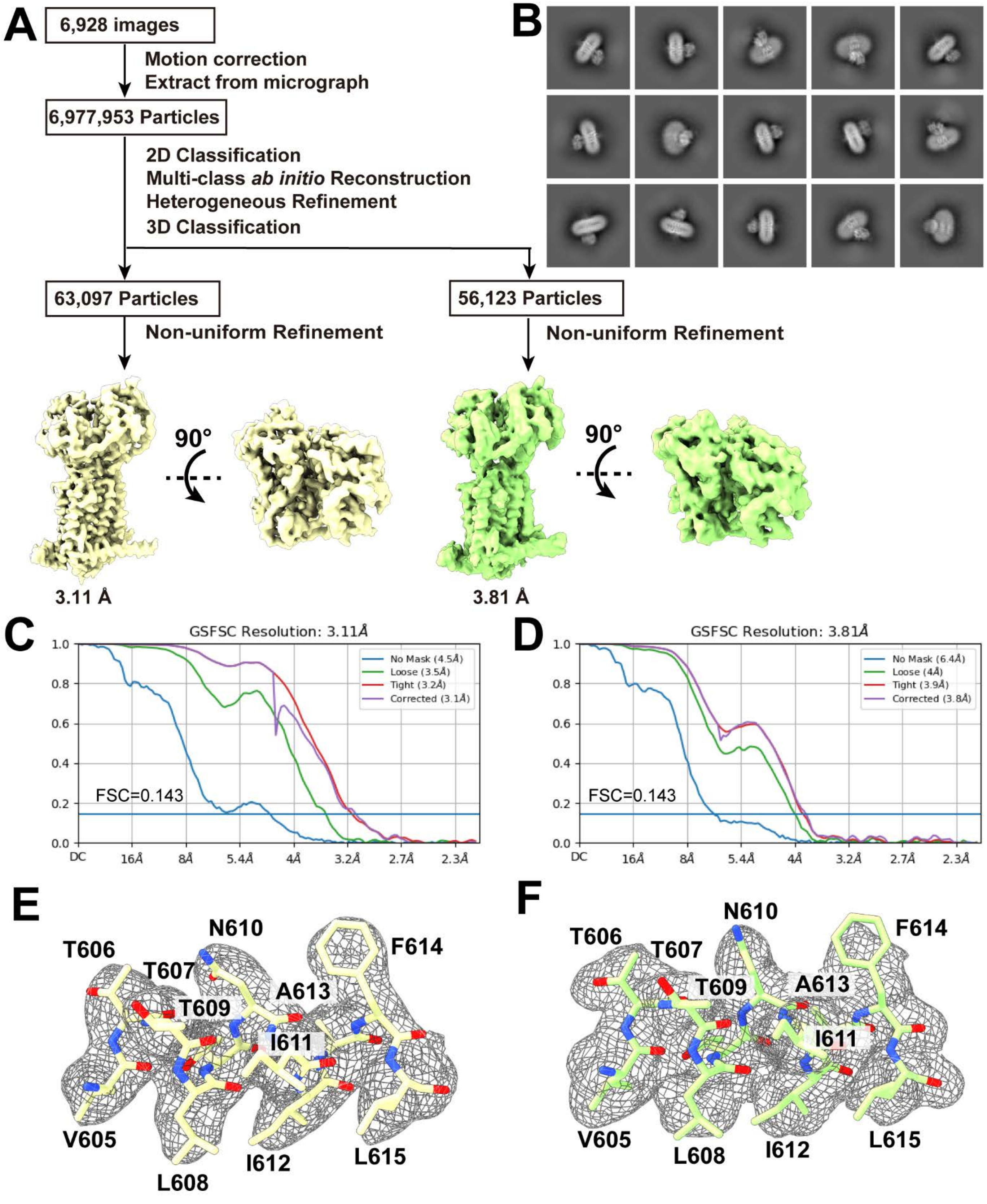


**Fig. S1. PfNCR1 data processing workflow.** (A) Data processing workflow of PfNCR1-I and PfNCR1-II. Side and top views of the PfNCR1-I and PfNCR1-II density maps. (B) Representative 2D classes of PfNCR1. (C, D) Gold-Standard Fourier shell correlation (GS-FSC) curves of PfNCR1-I and PfNCR1-II. (E, F) Local EM density maps of PfNCR1-I and PfNCR1-II.

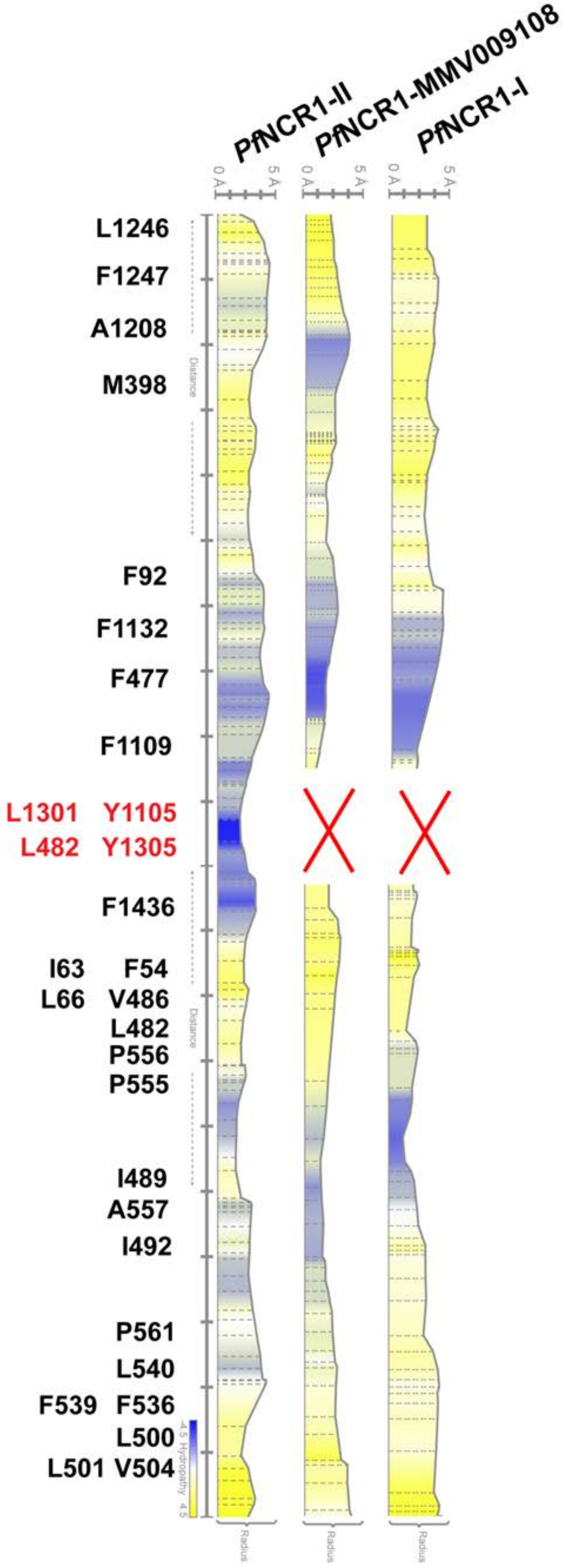


**Fig. S2. Tunnel formed by the PfNCR1 transporter.** This figure indicates calculated radii and important amino acids lining the tunnels formed by PfNCR1-I, PfNCR1-II and PfNCR1-MMV009108. The figure also indicates that the structure PfNCR1-II forms an open tunnel, whereas both the structures of PfNCR1-I and PfNCR1-MMV009108 depict closed tunnels.

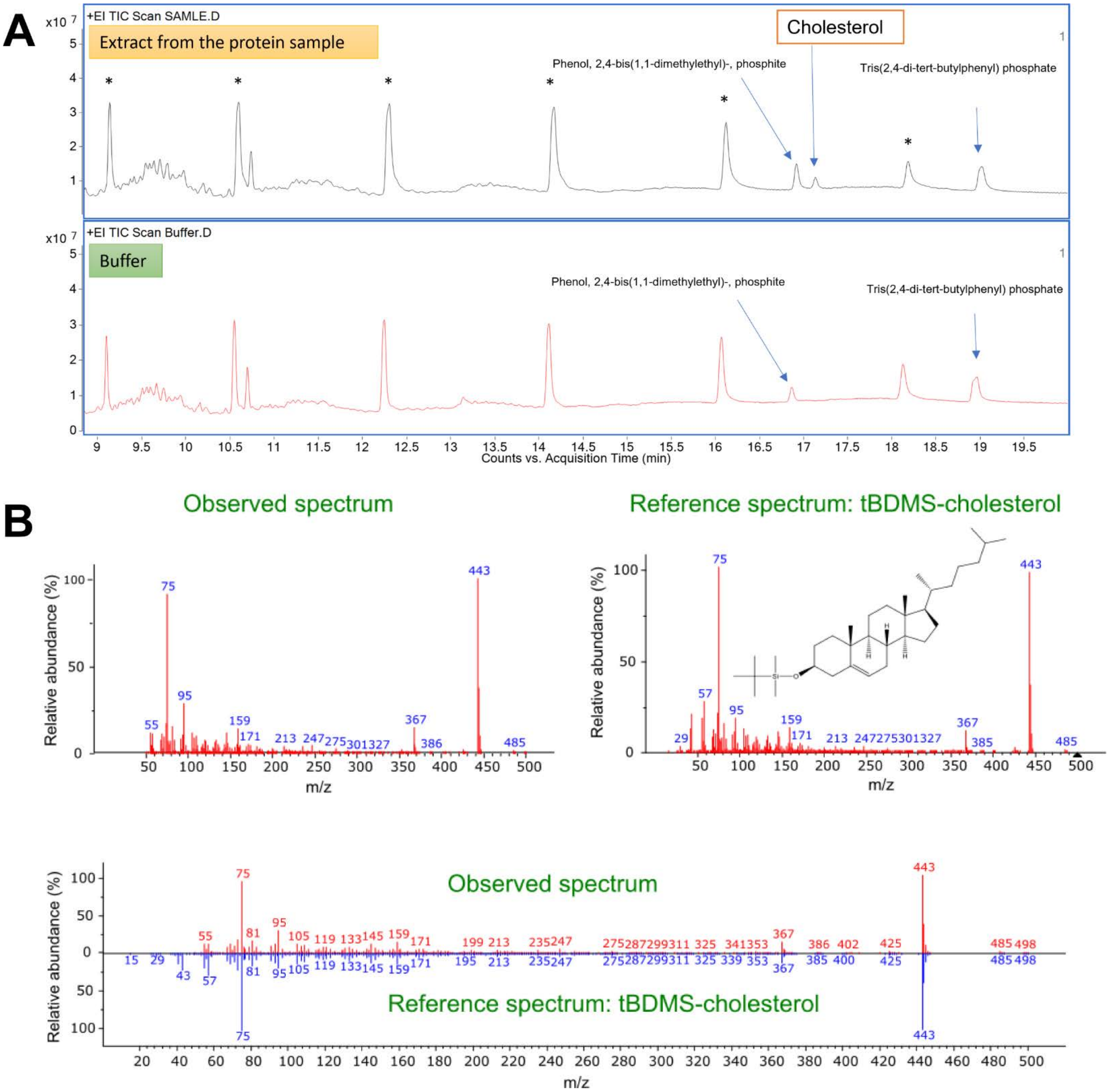


**Fig. S3. GC-MS spectra.** (A) GC-MS spectra of extract from the purified PfNCR1 protein sample (upper) and buffer of the sample (lower). (B) Comparasion of the observed and reference spectra. These spectra lead to the identification bound chlosterol in the PfNCR1 protein.

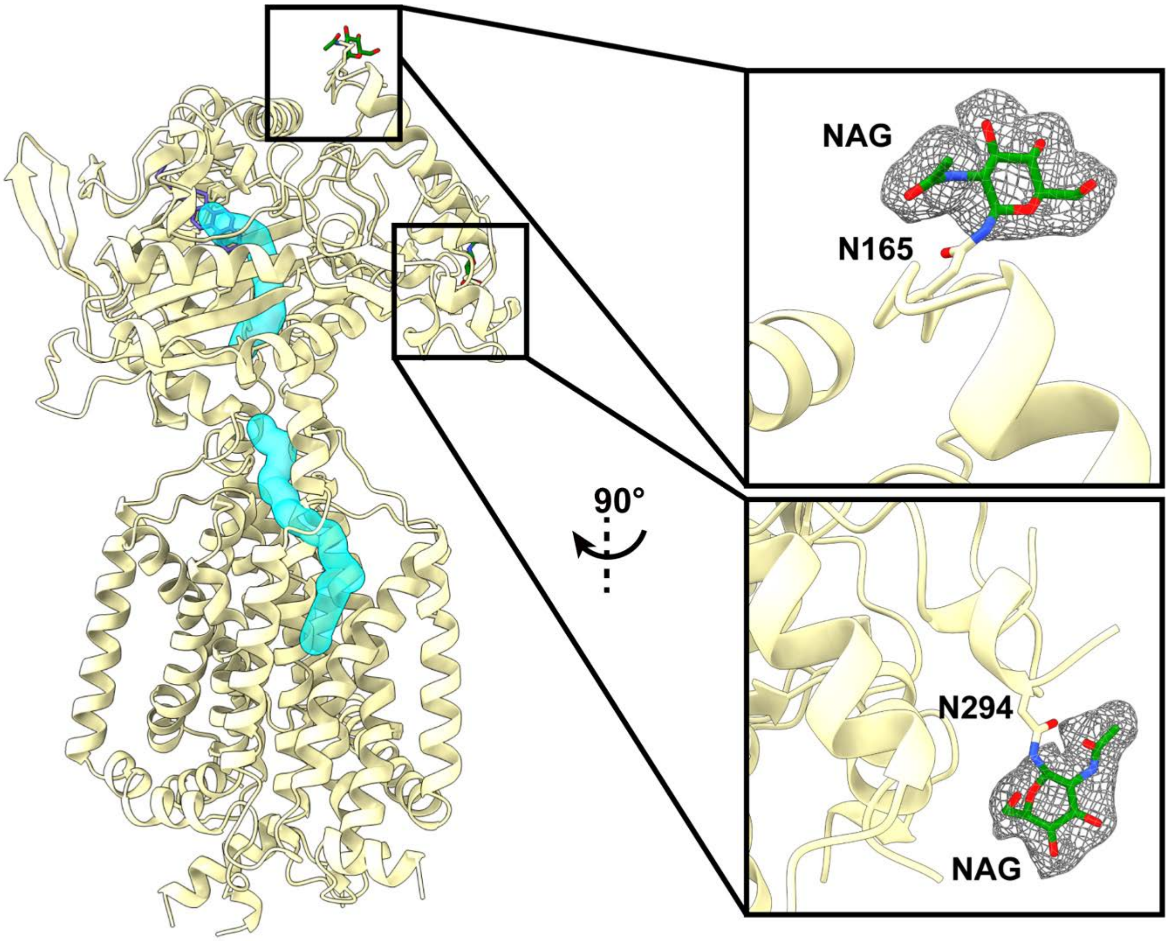


**Fig. S4. Cryo-EM densities of the two glycosylation sites of PfNCR1.** This figure includes thee secondary structural elements (yellow) of the PfNCR1 transporter. The tunnel formed by PfNCR1 is colored cyan. The cryo-EM densities showing the modifications of N165 and N294 are in gray meeshes. Based on the cryo-EM densities, each asparagine is connected to a NAG moiety.

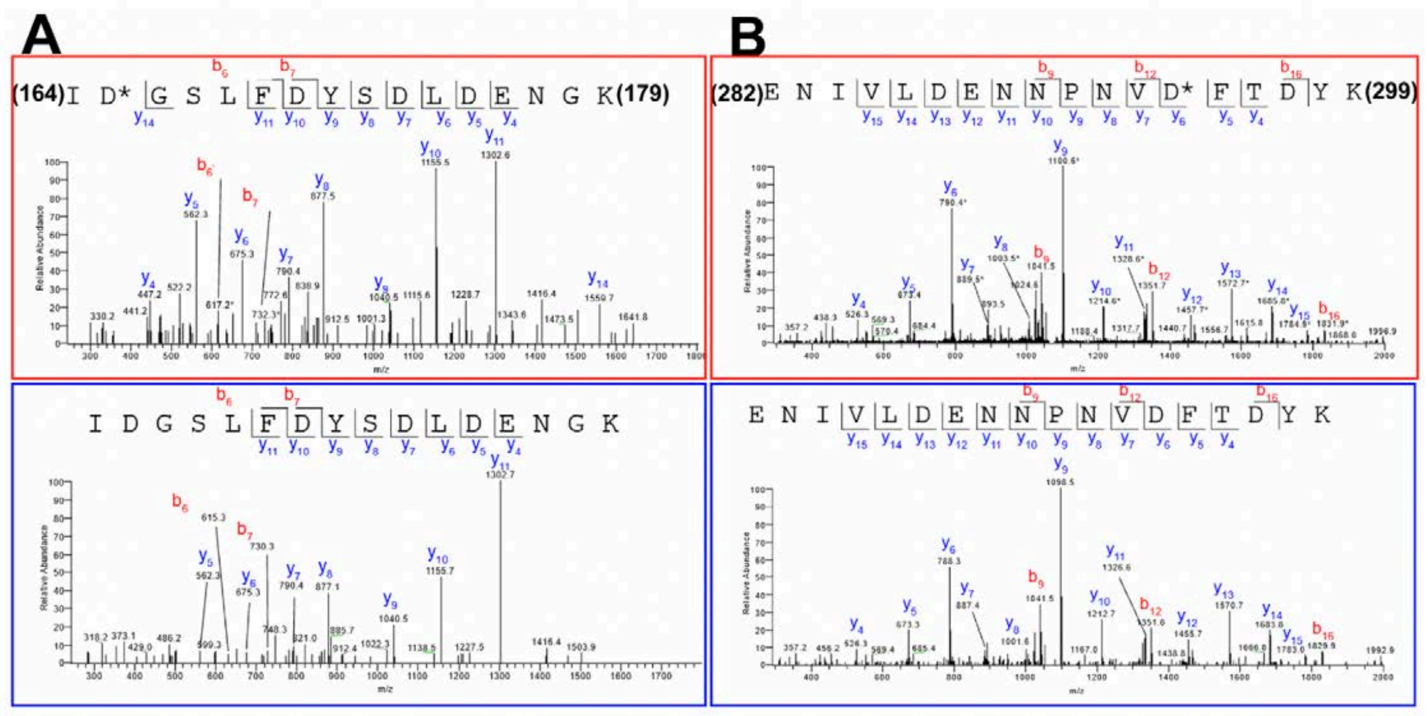


**Fig. S5. LC-MS-MS spectra of PfNCR1.** (A) Identified glycosylation site at residue N165. (B) Identified glycosylation site at residue N294. In both (A) and (B), the upper panel of the spectra are shown in the presence of $H_2{}^{18}O$, whereas the lower panel of the spectra are shown in the presence of $H_2{}^{16}O$. The amino acid sequence assigned from y-series ions is derived from the C-terminus of the peptide, and the amino acid sequence assigned from b-series ions is derived from the N-terminus of the peptide. The corresponding spectra of the glycopeptide treated with PNGase F are shown. The glycosylated asparagine residue has been converted to aspartic acid by enzymatic reaction. Its signal is shifted by +2 mass units (compared to the corresponding peptide incorporating $^{16}O$) as a result of $^{18}O$ incorporation.

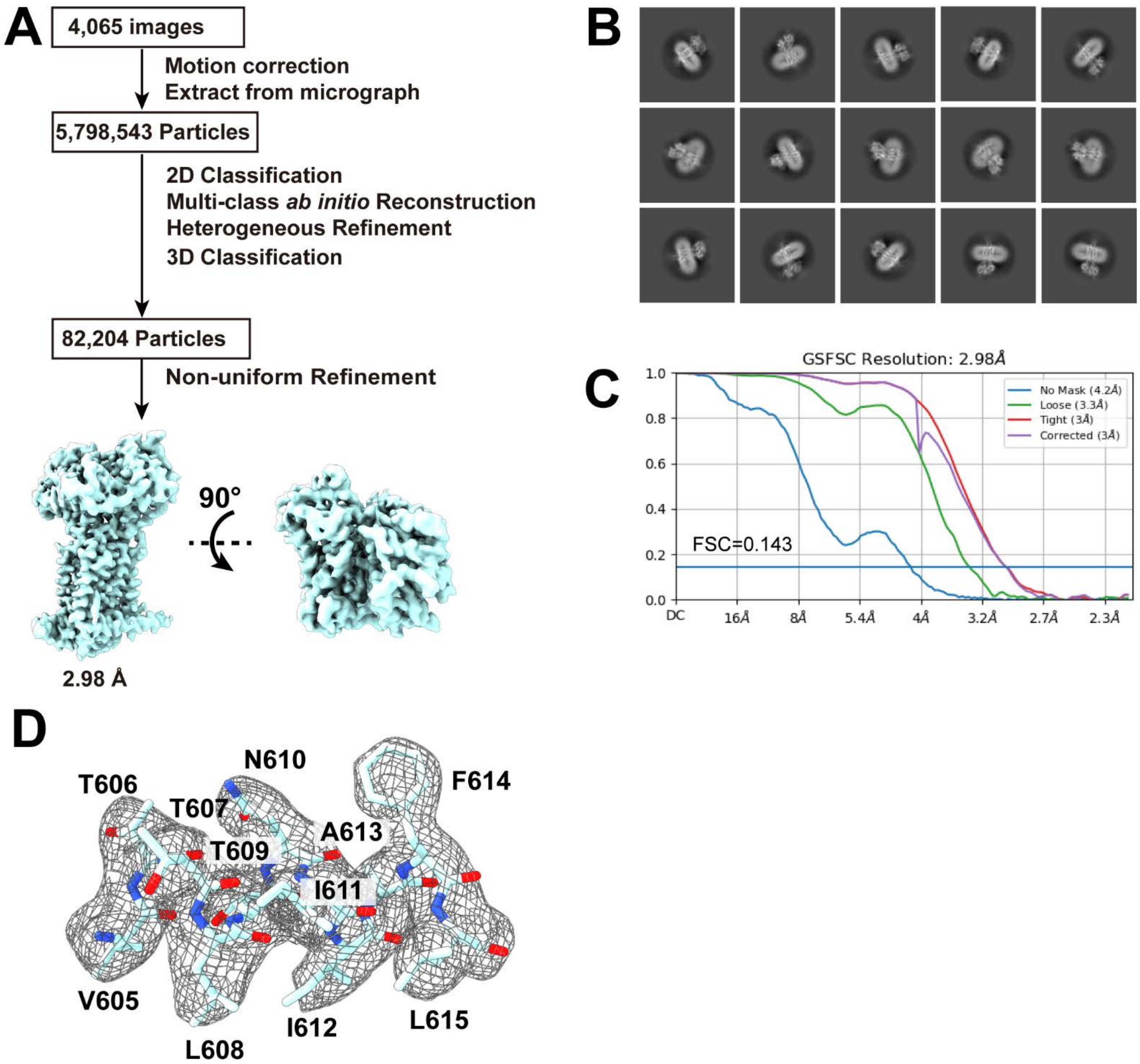


**Fig S6. PfNCR1-MMV009108 data processing workflow.** (A) Data processing workflow of PfNCR1-MMV009108. Side and top views of the PfNCR1-MMV009108 density map. (B) Representative 2D classes of PfNCR1. (C) Gold-Standard Fourier shell correlation (GS-FSC) curve of PfNCR1-MMV009108. (D) Local EM density map of PfNCR1-MMV009108.

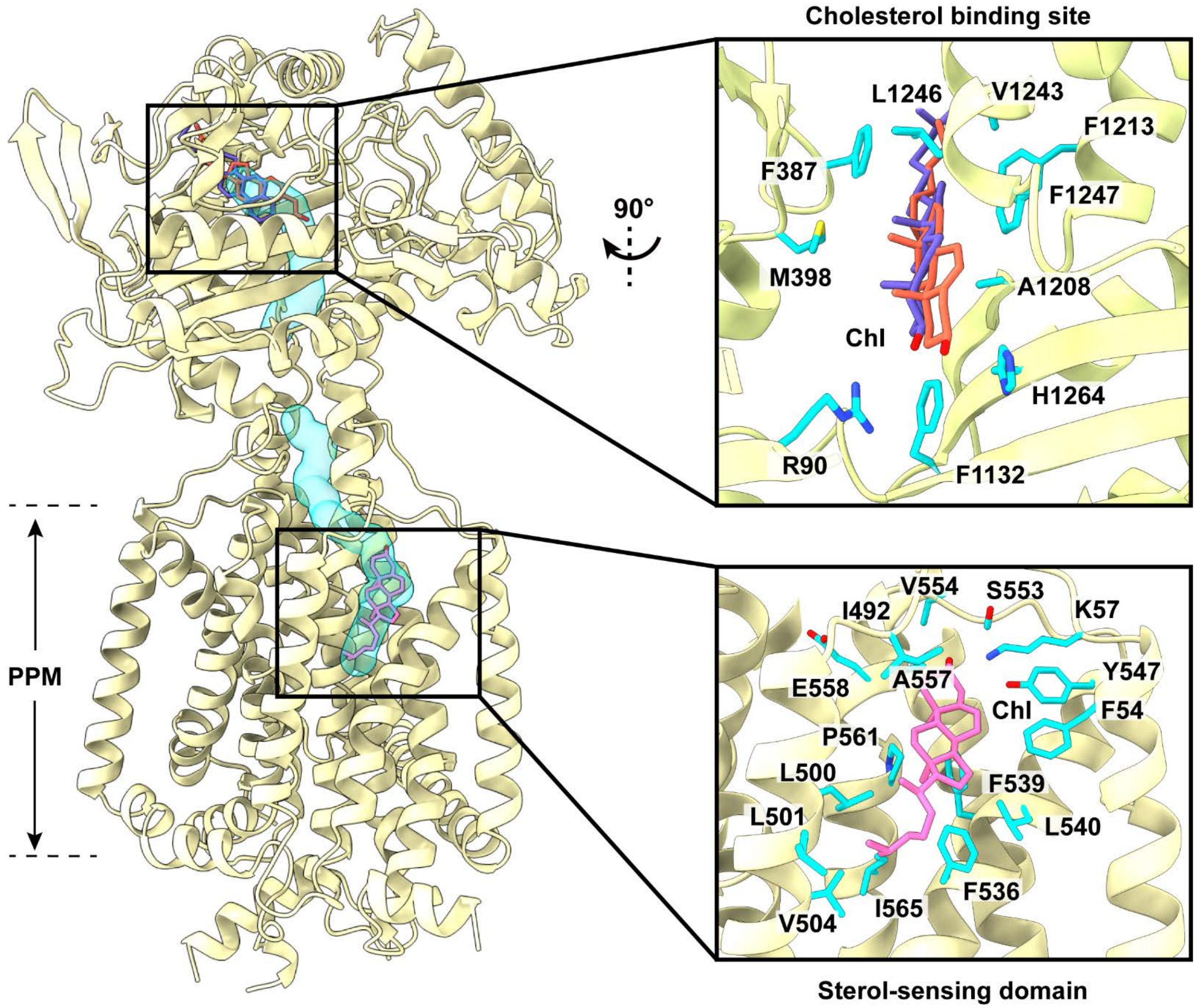


**Fig S7. Docking of Chl to the structure of PfNCR1.** The docking calculations indicate that there are two potent Chl binding sites located at the PV and PPM domains, respectively. Both docked Chl molecules are found within the tunnel (colored cyan) formed by the PfNCR1 transporter. The docked Chl molecule at Chl-binding site of the PV domain is shown as orange sticks. The bound Chl ligand identified from our cryo-EM structure is also included and shown as slate sticks. Residues responsible for Chl binding are in cyan sticks. The second predicted Chl-binding site is located at the SSD of the PPM domain of PfNCR1. The docked Chl molecule at this binding site is shown as magenta sticks. Residues predicted to be responsible for Chl binding are in cyan sticks.

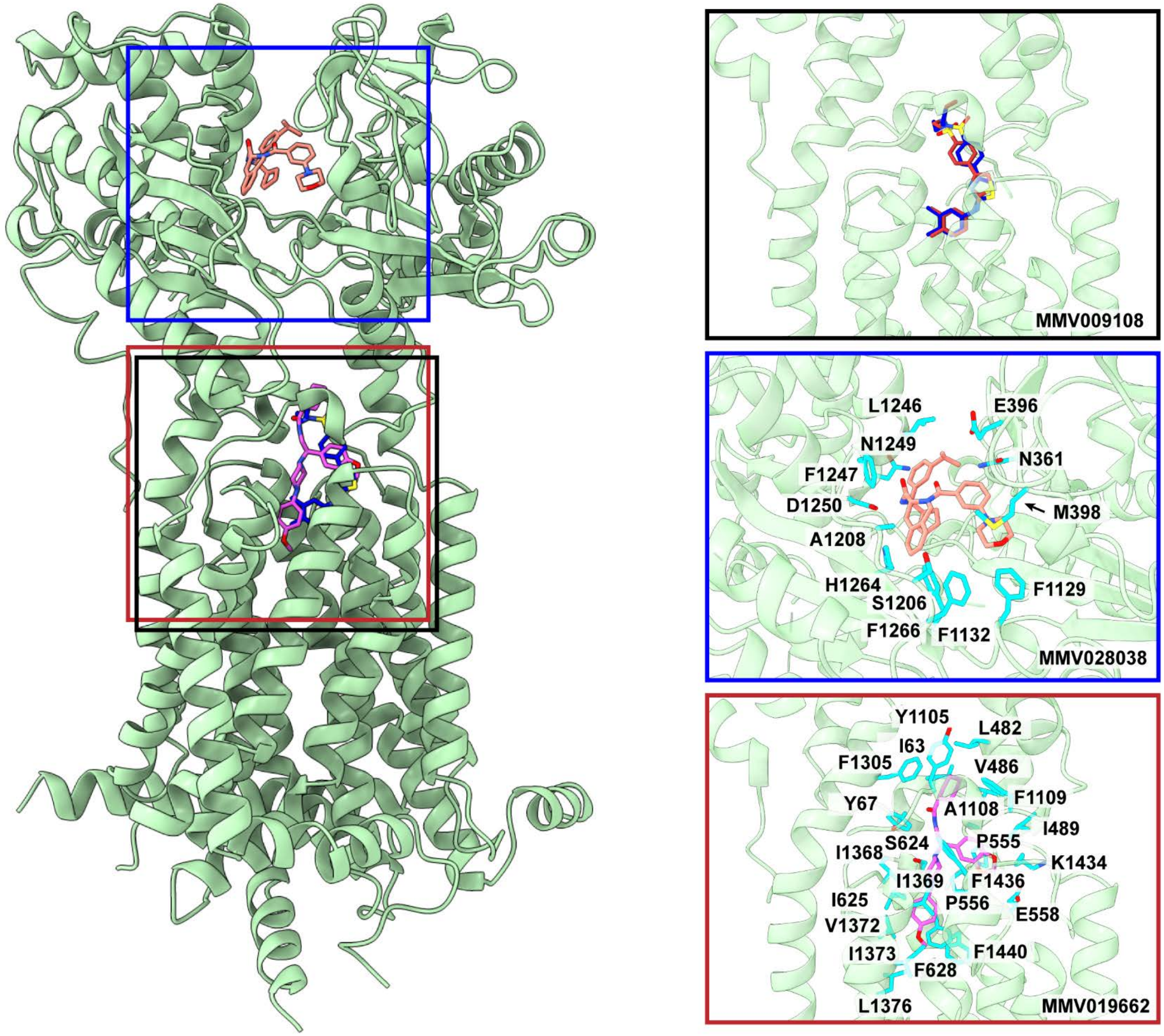


**Fig S8. Docking of inhibitors to the structure of PfNCR1.** The docking calculations indicate that both MMV009108 (blue) and MMV019662 (magenta) are bound in the middle of the transporter and nearby the restriction site of the tunnel formed by PfNCR1, whereas MMV028038 (orange) is bound within the cleft between subdomains PV1 and PV2, and overlapped with the Chl binding site at the PV domain identified by cryo-EM. The upper insert panel depicts bound MMV009108 (red) identified by cryo-EM and docked MMV009108 (blue) predicted via docking. The middle insert panel indicates the predicted MMV028038 binding site. The bound MMV028038 molecule is colored orange, whereas residues within 4 Å of bound MMV028038 are in cyan sticks. The lower insert panel shows the predicted MMV019662 binding site. The bound MMV019662 molecule is colored magenta, whereas residues within 4 Å of bound MMV019662 are in cyan sticks.

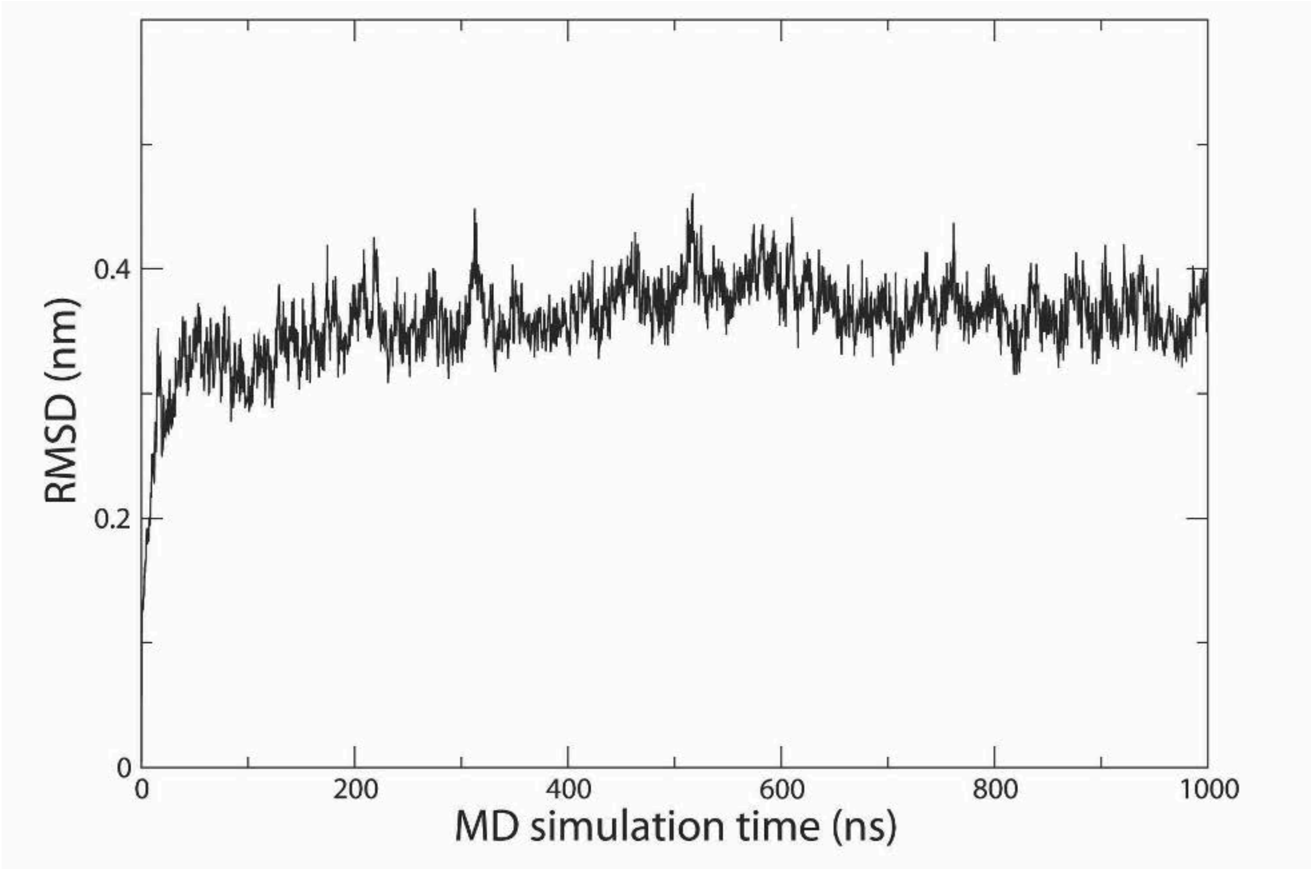


**Fig.S9. Results of 1 μs Simulation.** MD simulation results of the PfNCR1-I. The Cα atom RMSD (root mean square deviation) plot is based on the MD simulation trajectories (1 μs).

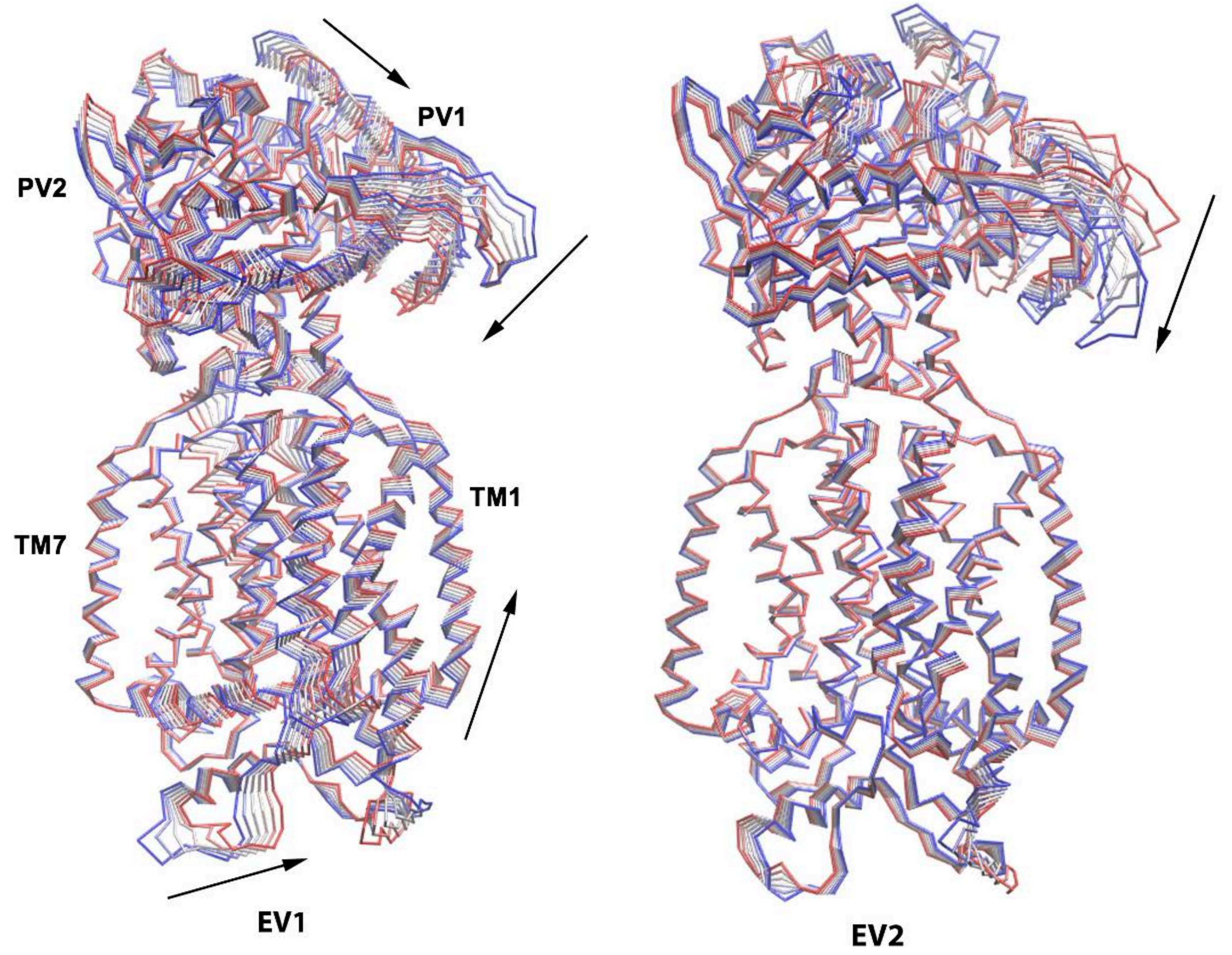


**Fig. S10. The first and second eigenvectors of PfNCR1 from PCA.** Black arrows show the rigid-body motions of PV and PPM domains. Six structures from each eigenvector were overlaid showing these rigid-body motions (blue to red).

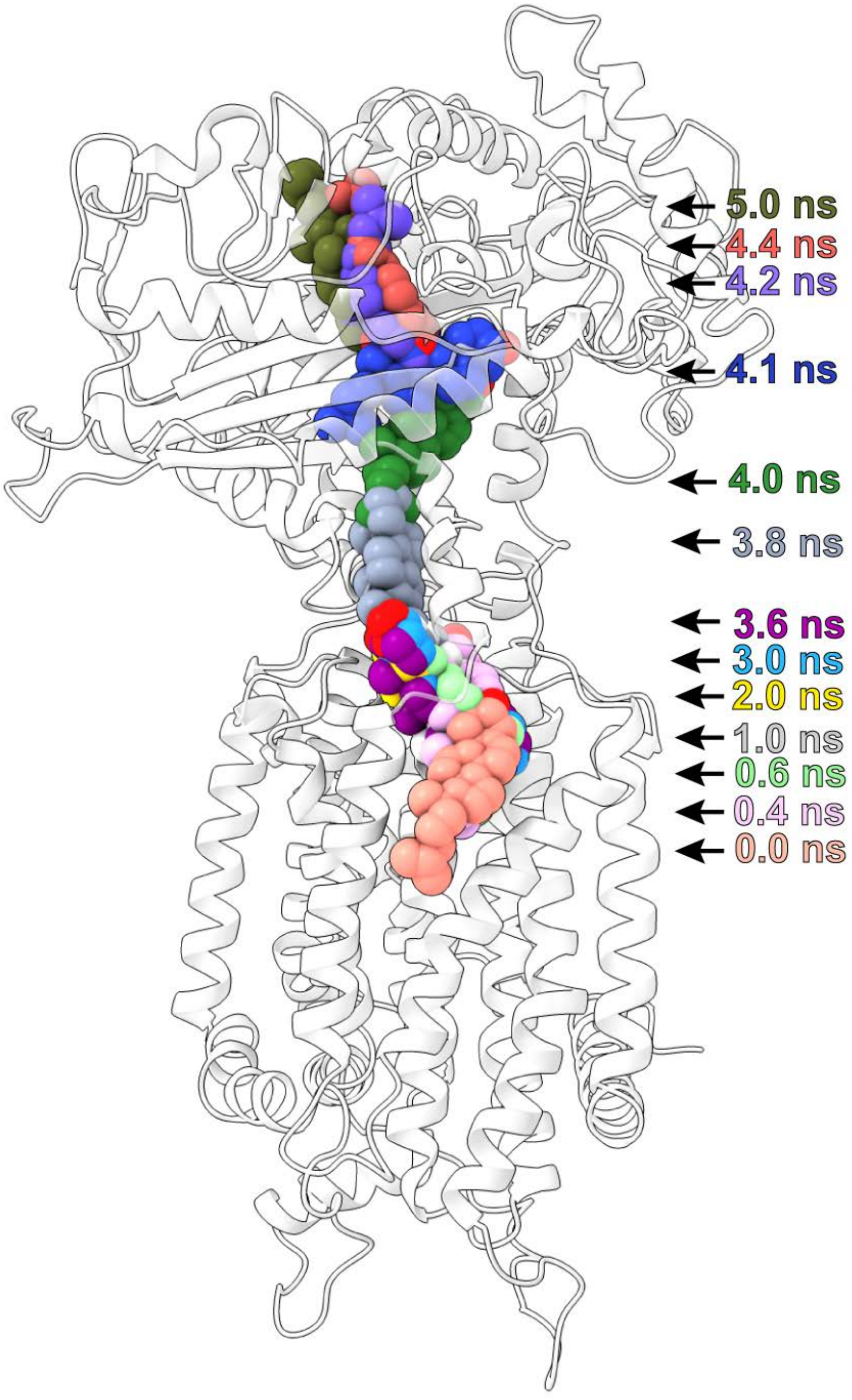


**Fig S11. Target MD simulations of the PfNCR1 transporter.** The calculations depict snapshots (0, 0.4, 0.6, 1.0, 2.0, 3.0, 3.6, 3.8, 4.0, 4.1, 4.2, 4.4 and 5.0 ns) of Chl shuttling across the tunnel formed by PfNCR1.

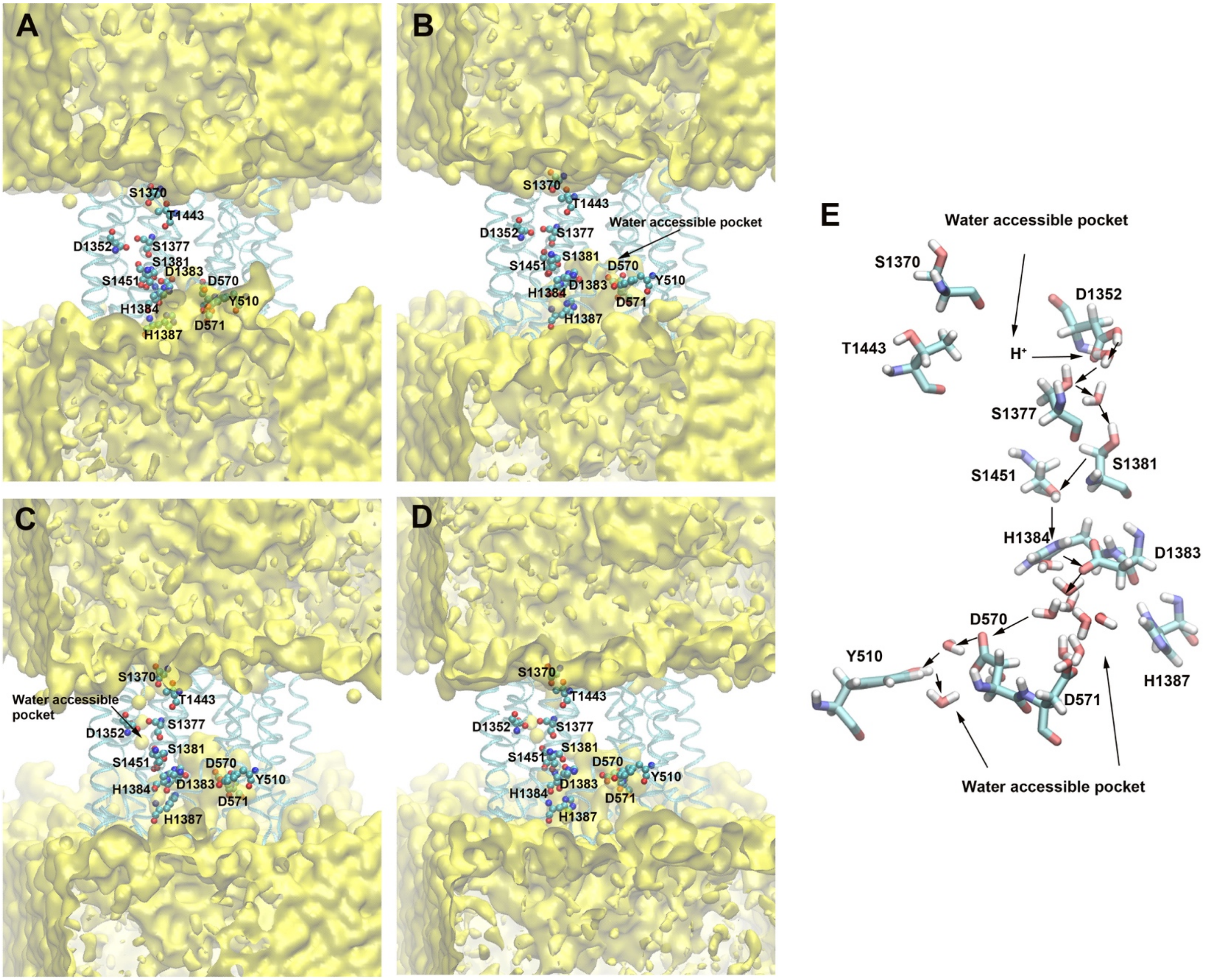


**Fig S12. Putative proton transfer pathway of PfNCR1.** This figure depicts snapshots of residues participating in the putative proton-relay network of PfNCR1 at (A) 0 ns, (B) 500 ns, (C) 593 ns and (D) 1000 ns. The simulations indicate that residues Y510, D570, D571, D1352, S1370, S1377, S1381, D1383, H1384, H1387, T1443 and S1451 form a proton-relay network to facilitate the transfer of proton across the membrane.

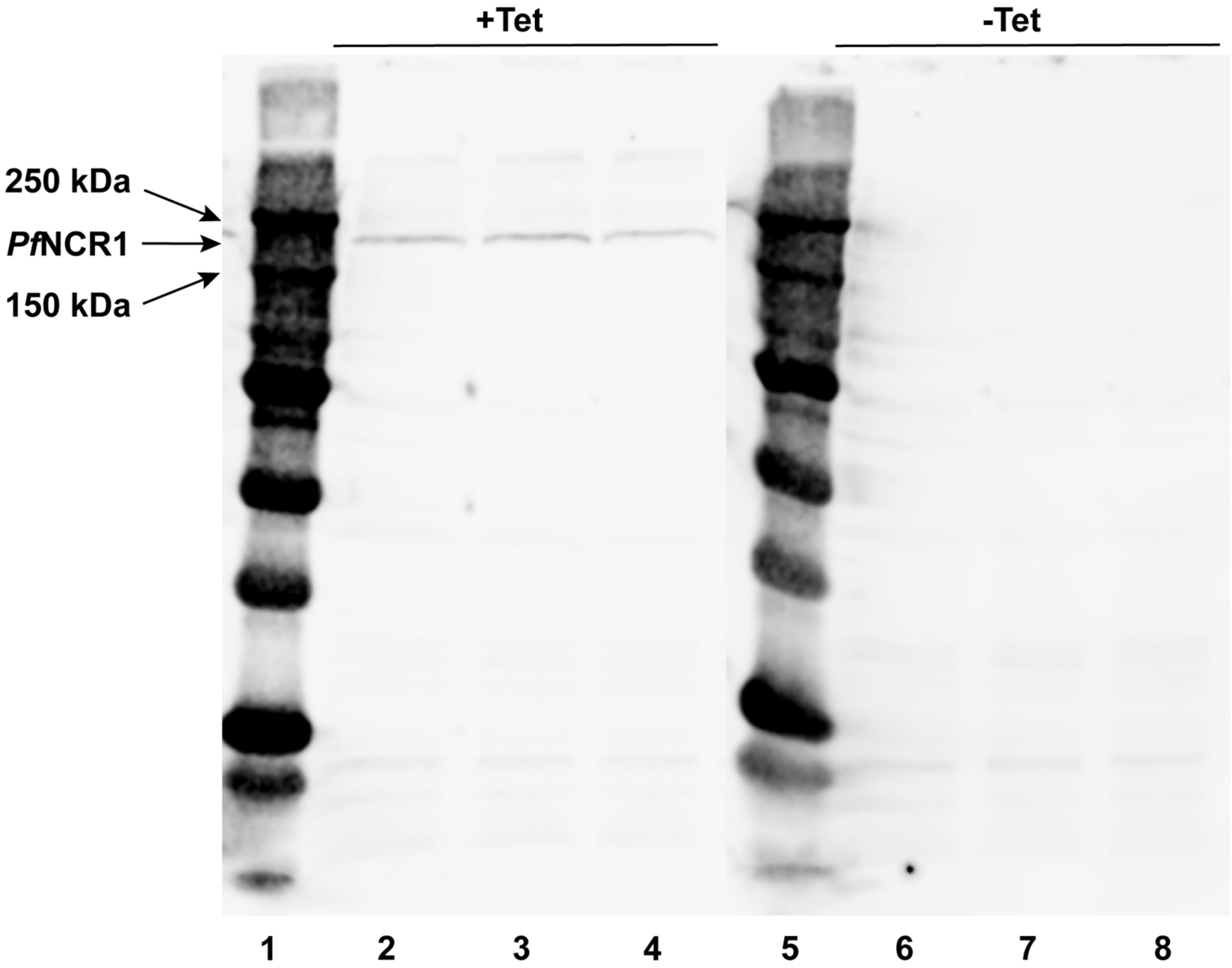


**Fig. S13. Expression of the PfNCR1 transporter.** This is an immunobolt against PfNCR1 of membrane extracts from 50 µg of dry HEK293 cells expressing the PfNCR1 transporter (lanes 1 to 8 from right to left: marker, HEK293 cells expressing PfNCR1 (+ tet), HEK293 cells expressing PfNCR1 (+ tet), HEK293 cells expressing PfNCR1 (+ tet), marker, control HEK293 cells (- tet), control HEK293 cells (- tet) , control HEK293 cells (- tet)).

**Table S1. PfNCR1 cryo-EM data collection and refinement statistics.**

| Data collection | PfNCR1 in the absence of MMV009108 | | | PfNCR1 in the presence of MMV009108 |
|---|---|---|---|---|
| Magnification | 81,000 | 81,000 | 81,000 | 81,000 |
| Voltage (kV) | 300 | 300 | 300 | 300 |
| Electron Microscope | Krios-GIF-K3 | Krios-GIF-K3 | Krios-GIF-K3 | Krios-GIF-K3 |
| Defocus (um) | -0.8 to -1.5 | -0.8 to -1.5 | -0.8 to -1.5 | -0.8 to -1.5 |
| Energy filter width (eV) | 20 | 20 | 20 | 20 |
| Pixel size (Å) | 1.07 (0.535) | 1.07 (0.535) | 1.07 (0.535) | 1.07 (0.535) |
| Total dose ($e^-/Å^2$) | 35.9 | 35.7 | 33.0 | 40.5 |
| Number of frames | 38 | 38 | 40 | 45 |
| Number of micrographs | 1,268 | 3,042 | 2,618 | 4,065 |
| Number of Initial particles | 6,977,953 | | | 5,798,543 |

| Refinement | PfNCR1-I | PfNCR1-II | PfNCR1-MMV009108 |
|---|---|---|---|
| Number of total particles | 63,097 | 56,123 | 82,204 |
| GS-FSC Resolution (0.143, Å) | 3.11 | 3.81 | 2.98 |
| <u>Model composition</u> | | | |
| Chains | 2 | 2 | 2 |
| Protein residues | 985 | 979 | 990 |
| Ligand | 3 | 2 | 4 |
| <u>r.m.s.d.</u> | | | |
| Bond lengths (Å) | 0.002 | 0.002 | 0.003 |
| Bond angles (°) | 0.460 | 0.480 | 0.445 |
| **Validation** | | | |
| MolProbity score | 1.51 | 1.89 | 1.30 |
| Clash score | 6.21 | 11.01 | 5.58 |
| <u>Ramachandran plot</u> | | | |
| Favored (%) | 98.87 | 98.55 | 98.37 |
| Allowed (%) | 1.13 | 1.45 | 1.63 |
| Disallowed (%) | 0.00 | 0.00 | 0.00 |
| CC Mask | 0.83 | 0.68 | 0.86 |
